# Herbivory can increase plant fitness via reduced interspecific competition – evidence from models and mesocosms

**DOI:** 10.1101/2024.04.20.589392

**Authors:** Laura Böttner, Fabio Dudenhausen, Sara Nouere, Antonino Malacrinò, Martin Schäfer, Joris M. Koene, Meret Huber, Shuqing Xu

**Affiliations:** Institute of Plant Biology and Biotechnology, University of Münster, 48143 Münster, Germany; Institute for Evolution and Biodiversity, University of Münster, 48149 Münster, Germany; Institute of Organismic and Molecular Evolution, Johannes Gutenberg University Mainz, 55128 Mainz, Germany; Department of Agriculture, Università degli Studi Mediterranea di Reggio Calabria, 89122 Reggio Calabria, Italy; Amsterdam Institute for Life and Environment, Section Ecology & Evolution, Vrije Universiteit, 1081 HV Amsterdam, Netherlands

**Author notes:** author for correspondence Meret Huber; Shuqing Xu. **<u>Statement of authorships</u>** conceptualization SX, MH; methodology SX, LB, FD, SN, MH; investigation LB, FD, SN, JMK, MS; visualization LB, AM, SX; funding acquisition SX, MH; project administration SX, MH; supervision SX, LB, MH; writing of the first draft of the manuscript LB, SX; all authors contributed substantially to revisions. Laura Böttner,; Fabio Dudenhausen,; Sara Nouere,; Antonino Malacrinò,; Joris M. Koene,; Martin Schäfer.

**Keywords:** duckweed (*Spirodela polyrhiza)*, algae, interspecific competition, intraspecific competition, plant-herbivore interactions, plant fitness

## Abstract

Herbivores are generally considered to reduce plant fitness. However, as in natural communities herbivores often feed on several competing plant species, herbivores can also increase plant fitness by reducing interspecific competition among plants. In this study, we developed a testable model to predict plant fitness in the presence of an interspecific competitor and an herbivore that feeds on both plant species. Our model allows to predict at which herbivore and competitor densities the focal species will benefit from herbivory. This can be estimated by quantifying the effects of the herbivore on the fitness of the focal plant and on its competitor, and by estimating the levels of intra- and interspecific competition in a pair-wise fashion, respectively. We subsequently validated the model in indoor microcosms using three interacting species: an aquatic macrophyte (the giant duckweed *Spirodela polyrhiza*), its native competitors (green algae), and its native herbivore (the pond snail *Lymnaea stagnalis*). Additional outdoor mesocosm experiments supported our model under natural conditions. Together, this study provides a conceptual framework to understand how herbivores shape plant fitness in a community context.

## Introduction

Plants are the backbone of most ecological interactions, and plant-herbivore interactions shape most of the planet’s ecosystems [1–3]. Herbivores are usually assumed to reduce plant fitness as herbivores consume plant material and thereby reduce growth and eventually reproduction (direct fitness costs). However, species do not occur in isolation but in a community, and herbivores often feed not only on a single but on multiple competing plant species. Thus, if an herbivore reduces the growth of a competitor, the herbivore may increase the fitness of other plant species, even if the plant is grazed upon (indirect fitness benefits through reduced competition). For example, molluscs may increase the growth of grasses (*Festuca rubra*) by decreasing the abundance of herbs [4]. Similarly, insect herbivores feeding on goldenrod populations indirectly led to a long-term increase in understory forb abundance by increasing light availability [5]. While these studies illustrate that indirect positive fitness effects of herbivores may outweigh their direct negative effects, we currently cannot predict the circumstances under which plants benefit from herbivores through reduced interspecific competition.

Predicting the net fitness effect of herbivores can be achieved by modifying and combining existing models of predator-prey interactions and two-species competition models [6–12]. Studies using mesocosms and microcosms can validate theoretic models; for example, [7]. Importantly, empirical studies should assess the consequences of herbivory and competition not only on plant phenotypes and growth, but also on plant fitness, that is, their contribution to the gene pool of the next generation [13]. However, this is challenging because many plants have long life cycles. Long-term herbivory exclusion experiments can overcome this limitation and thereby reveal the consequences of herbivory on plant abundance, community structure [14,15], diversity [4], and performance of alien plants under invasion [16]. However, in these long-term outdoor exclusion experiments, herbivores and competing plants continuously affect each other’s density, which prevents teasing apart, and therefore, quantifying the direct and indirect effects of herbivory. To test theoretic predictions when positive indirect effects outweigh the negative direct effects of herbivores, we need an experimental system in which we can manipulate the densities of both plant species.

One of the few experimental systems in which the densities of the plant, its herbivore, and its competitor can be easily manipulated is the duckweed-snail-algae system. Duckweeds are free-floating, aquatic plants of small body size and simple morphology: the green vegetative tissue forms a flat, thallus-like structure, the so-called frond. As most duckweed species, the giant duckweed, *Spirodela polyrhiza* (L.) Schleid, reproduces almost exclusively asexually by budding every 2-3 days under optimal conditions indoors [17,18] and every 3-7 days outdoors. Outdoors, the plant is frequently attacked by one of its major native herbivores, the holarctic common freshwater snail, *Lymnaea stagnalis*. This snail feeds not only on *S. polyrhiza* but also on green algae, such as the microalgae *Chlamydomonas* spp., which compete with *S. polyrhiza* for light and nutrients [19,20]. As the abundance of *S. polyrhiza*, its herbivore *L. stagnalis*, and its competitor *Chlamydomonas spp.* can be quantified and manipulated across multiple generations [21], these interacting species are ideal for testing the direct and indirect effects of herbivores on plant fitness.

Here, we first developed a theoretic model to understand the interaction between *S. polyrhiza,* its herbivore *L. stagnalis*, and its competitor *Chlamydomonas reinhardii*. Second, we used a reductionist setup for indoor model-guided assays to feed our theoretic model with data. Third, we performed indoor microcosm experiments as a proof of concept to validate our model, testing under which starting abundances of the competing species the presence of an herbivore promotes duckweed fitness by reducing interspecific competition. Fourth, we tested whether our findings hold true under outdoor conditions by quantifying the algal biomass and characterizing the algal community using amplicon metagenomics. Our findings highlight the importance of considering the real-world complexity when assessing the effects of herbivores on plant fitness.

## Material and Methods

### Model development and estimation of model parameters using laboratory microcosms

First, we built a generic model that incorporates both plant-herbivore interactions and plant-plant competition based on existing species-competition models [6–8,10–12,22].

For simplicity, we focused on two competing species that can be attacked by the same herbivore. The population growth of the two plant species, which is affected by competition (intra- and interspecies) and herbivory, can be expressed as

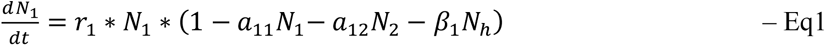

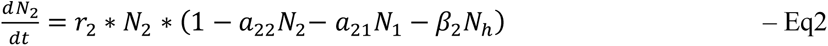

where the *r*_1_ and *r*_2_ are the intrinsic growth rates of the two species. *N*_1_ and *N*_2_ are the numbers of individuals from the two species at a given time, *α*_11_ and *α*_22_ refer to intraspecific competition. *α*_12_ and *α*_21_ are interspecific competition coefficients describing the effects of the species associated with the second number in the subscript on the species associated with the first number in the subscript. *β*_1_ and *β*_2_ refer to herbivore consumption rates of species 1 and 2, respectively. *N*_ℎ_ refers to herbivore abundance. For simplicity, we assumed a constant herbivore population size.

To feed our theoretic model with experimental data, we used model-guided laboratory two-species assays, which allowed us to estimate the model coefficients. As model species we used *S. polyrhiza* (genotype 9500, originating from Germany), *C. reinhardii* (initial culture originating from a natural pond in Münster, Germany), and simultaneously hermaphroditic pond snail *L. stagnalis* (initial population originating from a long-standing laboratory culture from the Vrije Universiteit Amsterdam, Netherlands, [23]). Precultivation and indoor assays were carried out in a growth chamber at 26 °C with a light intensity of 135 µmol m^-2^s^-1^ under long-day conditions (16h:8h). All species were first cultivated in individually optimized media [24] (SI Table S1) and then preadapted to common experimental conditions (SI Table S2, SI Text S1). The snails were starved for 24 h under experimental conditions prior to the assays. For indoor assays, we used transparent plastic beakers (diameter 11 cm, PP) with 150 m L of Mix-Medium (SI Table S2). The beakers were covered with dark foil to avoid the incidence of light, and beakers were covered with transparent and perforated plastic lids to ensure gas exchange but to prevent evaporation and escape of herbivores.

To quantify *C. reinhardii*, we generated a reference curve using a Thoma cell counting Chamber and measured the corresponding optical density with a plate reader (Tecan infinite 200 with the software MagellanPro v7.3 2019) for OD_750 nm_ [21] with pure Mix-Medium as a blank, covering a *C. reinhardii* concentration range from 0 to 22 [10^6^ cells mL^-1^] (SI Fig. S1). To estimate the intra- (α_11_) and interspecies (α_12_) coefficients of *S. polyrhiza* of our theoretic model, we quantified fitness, measured in terms of the number of individuals (expressed as per capita growth) of *S. polyrhiza* when competing against varying densities of itself, as well as when grown in the presence of a constant concentration of *C. reinhardii* as a competitor. We grew duckweed with a starting population size of 5, 50, 100, 300, 600, or 1000 fronds in triplicate in the absence and presence of a fixed algae concentration of 1.33 [10^6^ cells mL^-1^]. We reset the algae concentration to 1.33 [10^6^ cells mL^-1^] daily for seven consecutive days, using centrifugation. To maintain the same medium with respective nutrient level changes throughout the course of the experiment, we reused the supernatant medium of each individual experimental beaker and added newly cultivated algae cells (daily preadapted to Mix-Medium) at the respective concentrations. To quantify *S. polyrhiza* on day seven of the experiment, we counted the number of live fronds, referring to green fronds with intact reproductive pouches. All the visible small daughter fronds were counted as individuals. Per capita growth per day is expressed using the formula ln(1+((number of fronds at day 7 – number of fronds at day 0) / number of fronds at day 0 / 7 days)).

To estimate the coefficients α_22_ and α_21_ describing the intra- and interspecific coefficients of *C. reinhardii*, we grew algae in defined starting concentrations (0.0095, 0.59, 0.13, 1.64, 3.91, 7.66, 9.24 [10^6^ cells mL^-1^]) in the absence (n=4 per gradient step) and presence (n=4 per gradient step) of 100 fronds. To keep the number of fronds constant across seven days, we manually reduced duckweed to 100 individuals on days two and five of the experiment. Per capita growth was calculated analogously to the per capita growth of *S. polyrhiza*. Here, we assume a linear relationship for the interspecific competition.

To determine the individual feeding rate of *S. polyrhiza* (β_1_) by a snail within 24 h, we subjected 100 fronds to one snail (n=6) or none as a control (n=4). We harvested all remaining fronds after 24 h. Similarly, to determine the individual feeding rate of *C. reinhardii* (β_2_) by a snail within 24 h, we started with a concentration of 12.6 [10^6^ cells mL^-1^] and one snail (n=8) or none as control (n=4). We again measured the OD after 24 h to determine the remaining algae concentration. For all consumption rates we assumed a constant feeding rate within 24 h. The data from the experiments described above allowed us to estimate the competition coefficients (α_11_, α_22_, α_12,_ α_21_) and consumption coefficients (β_1_, β_2_) based on our proposed model equations using R [25] (see the SI R Markdown script). Data were visualized using *ggplot2* [26].

The 95% confidence intervals of each parameter were estimated using a bootstrapping approach with 500 repetitions. Based on the extracted coefficients, we were able to test the predicted scenarios from the SI Fig. S2 to determine the scenario that best describes the duckweed-algae-snail interaction.

### Testing our theoretic model within laboratory microcosms

We tested within laboratory three-species microcosms under which starting conditions *S. polyrhiza* benefitted from the presence of a snail herbivore when competing with *C. reinhardii*. We included a treatment with a high number of duckweeds competing against a low concentration of algae, and vice versa, as well as two treatments with intermediate densities. We grew these combinations in the presence or absence of none, one, or two snails. We excluded replicates in which the populations collapsed, or snails died within 14 days from the analysis, resulting in a data set of n=3-8 per treatment. The details can be found in the Supporting Information (SI Text S2). On day 14 of the experiment, we counted the number of live fronds, measured the OD of algae, and calculated the per capita growth of both competing species. Data were analysed using Wilcoxon tests to compare per capita growth between the herbivory and control treatments.

### Experimental validations in the field

To test our model in a natural setting, we set up an outdoor experiment in 2019 in Münster, Germany (51°57’54.0’N, 7°36’22.4‘E), in which *S. polyrhiza* was exposed to a natural community of green algae in the presence and absence of snail herbivory. Eight experimental ponds (60 × 80 × 32 cm; AuerPackaging, Amerang, Germany) were buried on the rim of the soil and filled on June 26^th^, 2019 with 120 L of tap water, 6 L of commercial pond soil (∼ 100 mg/L each of organic N, P_2_ O_5_, K_2_ O, ∼160 mg/L Mg, pH (CaCl_2_)= 4.2, salts 1.0 g/L; Floragard, Oldenburg, Germany), and 40 mL of commercial liquid fertilizer (organic NPK 3,1+0,5+4,1; COMPO BIO, Compo GmbH, Münster, Germany) based on pretests allowing sufficient growth similar to natural pond systems. To allow the establishment of a natural (phyto-)plankton community, ponds were inoculated with filtered water (via coffee filter) from three natural ponds located within and around Münster Municipality, where duckweeds (*S. polyrhiza* and *Lemna sp.*) naturally occur (SI Table S3). The experimental ponds were covered with a stainless-steel mesh (mesh opening 0.63 mm, wire diameter, 0.224 mm; Haver&Boeker, Oelde, Germany) to avoid debris and small animals from entering the ponds. The ponds were divided into two compartments by using the same metal net. Each compartment received 1000 *S. polyrhiza* fronds that were pre-cultivated indoors (S1 Text 1). The plants were shaded with a shading net during the first week to facilitate plant establishment. Then, on July 15th, the experiment began with eight snails (shell size 14-32 mm), which were added to one of the compartments of each pond; the other compartment served as a control. As the two compartments were separated with a fine metal net, water and dissolved nutrients, but not herbivores, could be exchanged between compartments. To compensate for the water loss due to evaporation, the ponds were refilled with tap water once a week. In week six, aphids invaded three ponds and extinguished duckweed populations. To avoid further contamination, we removed infested ponds from the field and excluded all the data from our analysis. Therefore, all the data presented correspond to n=5 per treatment until week nine, which was the end of the experiment.

To track the growth of the duckweed populations, we took pictures of the populations weekly and estimated the surface area covered by plants over time using the ImageJ software[27]. We further tested whether herbivory altered the biomass of fronds by sampling 20 randomly selected fronds from each treatment zone and pond at weeks 0, 2, 3, 4, 5, 7, 8, and 9. Further details and statistics can be found in SI Text S3. To test whether herbivory affects metabolite levels of *S. polyrhiza*, we harvested fronds at weeks 0, 3 and 6, and analysed a wide set of primary and specialized metabolites via HPLC-UV/VIS and LC-MS. The details can be found in SI Text S4. To test whether water exchange through the net maintained equal nutrient levels between the herbivore and control compartments, we analysed the nutrients in each treatment zone per pond at weeks 0, 3, 5, 7, and 9 via ion chromatography (792 basic IC, 732 detector, Metrohm, Herisau, Switzerland). Details on the water sampling and analysis can be found in SI Text S5.

We also collected algal samples from the field. We first identified the emerging algal species based on their morphology. We characterized free-swimming microalgae emerging within the first half of the season as well as sediment-rooted macroalgae that emerged during the second half of the season. To test whether herbivory alters the biomass of sediment-rooted macroalgae, we harvested all sediment-rooted macroalgal materials at the end of the season. The harvested macroalgae material was washed several times in tap water, frozen at -20 °C, and freeze-dried. Dry weight was determined and analysed using a linear mixed effect model with *treatment* (herbivory/control) as a fixed factor and *pond* as a random factor, using the formula: ∼ treatment * (1|pond).

To test whether algae and associated (cyano-)bacterial communities differ under herbivory and control conditions outdoors, we used an amplicon metagenomic approach. We selected two types of samples: First, we sampled 30 fronds of each treatment and control pond (n=5) at weeks 0, 4, and 7 to characterize the free-swimming microalgae (18S) and (cyano-)bacteria (16S) communities that were attached to or growing as endophytes of *S. polyrhiza* fronds. Upon harvest, the fronds were washed with ultrapure water and frozen at -20 °C. Secondly, we harvested macroalgal material with adherent microalgae at the end of the season, as described above, to characterize the (macro-)algal community (18S) of each treatment and control pond (n=4-5, as one herbivore compartment did not contain any macroalgae).

Freeze-dried fronds were ground with steel balls inside a tissue lyser (1.5 min at 30 Hz). DNA was extracted using a phenol-chloroform protocol. Samples were then shipped to Novogene Ltd. for amplicon metagenomic library preparation and sequenced on an Illumina NovaSeq 6000 S4 250PE flow cell. (Cyano-) Bacterial communities were assessed by amplifying the V4 region of the 16S rRNA (primers 515F and 806R), whereas the eukaryotic community (including algae) was determined by amplifying the V4 region of the 18S rRNA (primers 528F and 706R).

Macroalgal material with adherent microalgae was freeze-dried, weighed, and ground using a mortar and pestle. DNA was extracted from approximately 30 mg of freeze-dried ground tissue using the Plant II Mini Kit (MacheryNagel, Düren, Germany) according to the manufacturer’s instructions with the following adjustments: 400 µL P1 (CTAB-based), lysis for 1 h at 65 °C, and elution for 20 min at 65 °C. The samples were then subjected to StarSeq (Mainz, Germany) for library preparation (18S, primers 528F, and 706R) and sequencing using Illumina MiSeq (300PE).

Raw data were processed using *TrimGalore* v0.6.7 to remove Illumina adaptors and discard low-quality reads. Paired-end reads were processed using the *DADA2* v1.22 [28] pipeline implemented in R to remove low-quality data, identify ASVs, and remove 10 chimeras. Taxonomy was assigned using the *SILVA* v138 database [29] for 16S data or the *PR2* v4.14 database [30] for 18S data. Data were analysed using R v4.1.2, with the packages *phyloseq* v1.38 [31], *vegan* v2.6 [32], *DESeq2* 1.34 [33], and *lme4* 1.1.33 [34]. Analysis of 18S was restricted to the phylum “Archaeplastida,” which includes alga. Before downstream analyses, contaminants were removed using the decontam v1.14 [35] and data from non-template control libraries. We also removed reads identified as “chloroplast” or “mitochondria”, and singletons.

Distances between pairs of samples in terms of community composition were calculated using an unweighted UniFrac matrix and then visualized using a nonmetric multidimensional scaling (NMDS) procedure. Differences between sample groups were inferred through permutational multivariate analysis of variance (PERMANOVA, 999 permutations), specifying treatment (herbivory/control) as fixed factor, and using the factor “*pond*” to stratify permutations. Differences in microbial diversity between groups were assessed by calculating the Shannon diversity index and by fitting a linear mixed effects model using the formula Shannon _index ∼ treatment * timepoint * (1|pond). ASVs that were differentially abundant between treatments (herbivory/control) were identified using *DESeq2* with FDR-corrected *P*<0.05.

## Results

### A model for plant fitness consequences under herbivory in the presence of a competitor

We proposed a testable theoretic model describing the interaction of a focal species 1, its competing species 2 and an herbivore feeding on both species (meta-model Fig. 1a). The model aimed to predict under which starting abundances of each species, our focal species will benefit under herbivory by reducing interspecific competition. Based on the Eq1 and Eq2, we derived the isoclines of two species, and the lines that represent the zero-growth rate of each species can be expressed as:

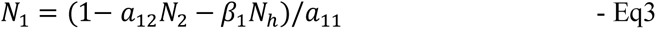

and

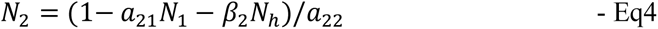

**Figure 1.**
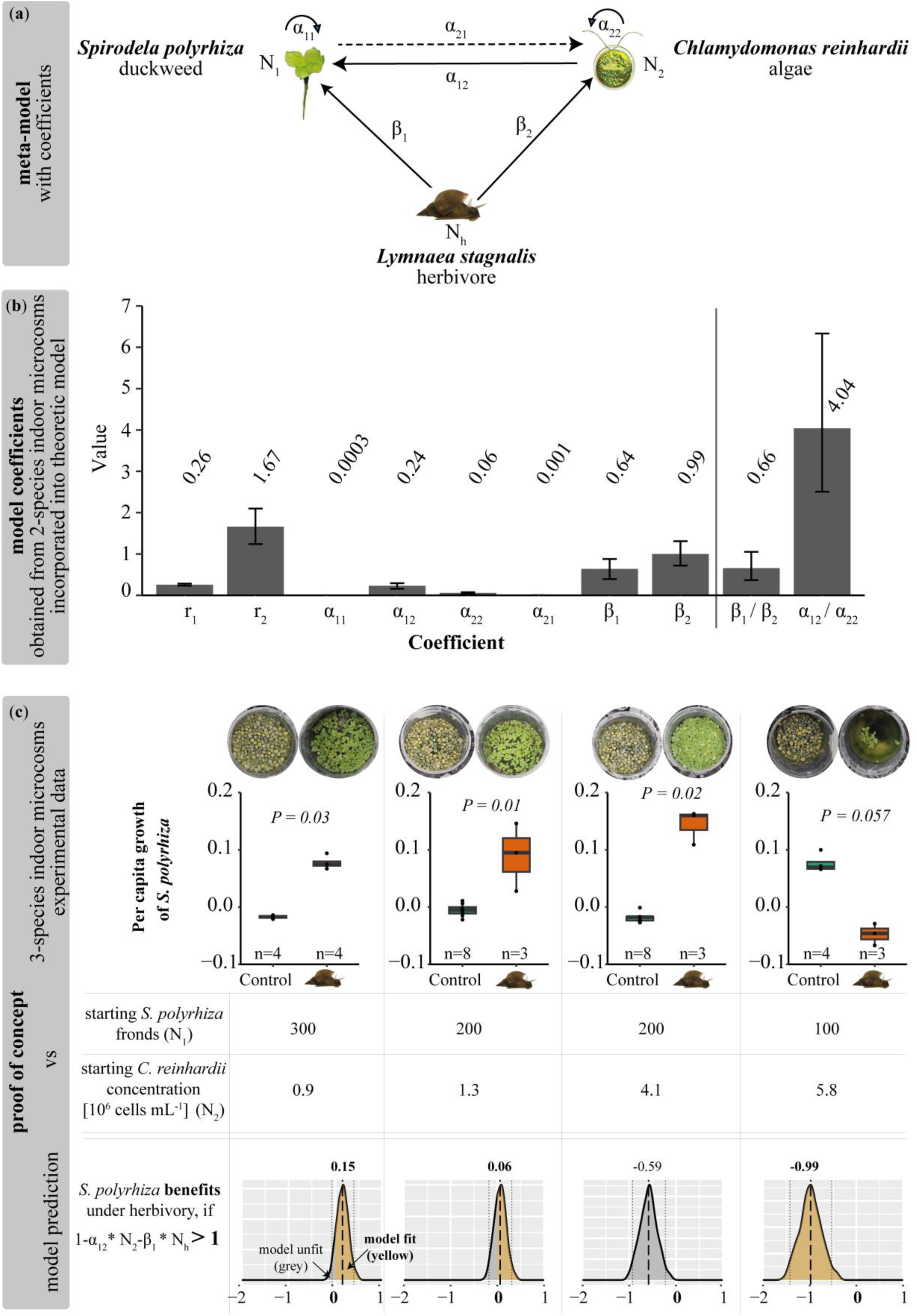
Modelling approach and experimental proof of concept to test under which starting conditions duckweed benefits from herbivory when competing with algae. (a) Meta-model representing the causal relationships among our focal plant species *Spirodela polyrhiza*, the green microalgae *Chlamydomonas reinhardii* and the herbivorous snail *Lymnaea stagnalis*. Both competing species underly intraspecific (α_11_, α_22_) and interspecific (α_12_, α_21_) competition and a fixed abundance of the herbivore (*N*_ℎ_) has individual consumption rates of each organism (β_1_, β_2_). (b) Estimated coefficients based on two-species laboratory assays (mean and 95 % confidence interval). Numbers above bars give mean values. Based on our model, species 1 benefits under herbivory if 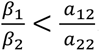. (c) Comparison between three-species indoor-microcosm data and predicted outcome using our model approach. Starting abundances of interaction partners determined if *S. polyrhiza* benefitted under herbivory in the presence of *C. reinhardii* as competitor. *P*-values refer to Wilcoxon tests, n=3-8. The prior conditions (1 − *α*_12_ ∗ *N*_2_ ∗ *β*_1_ ∗ *N*_ℎ_ > 1) in which snail herbivory benefits duckweed are shown at the bottom. Plots of model predictions indicate means (values on top, black dashed lines) and 95 % confidence intervals (grey dashed lines) estimated based on 500 bootstraps. Yellow shaded areas indicate congruence of theoretic model (1 − *α*_12_ ∗ *N*_2_ ∗ *β*_1_ ∗ *N*_ℎ_) and experimental outcome (values > 0 indicate a fitness benefit for *S. polyrhiza*). Picture of *C. reinhardii* was created with Biorender.com.

A graphic representation of the isoclines is shown in SI Figure S2. When the two species do not coexist, herbivory negatively affects plant fitness. If the two species coexist (*N*_1_ > 0 and *N*_2_ > 0), they must share one or more points on their isoclines, which can be calculated by substituting Eq3 into Eq4 and vice versa, as follows:

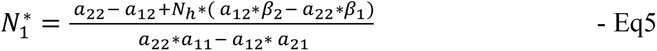

and

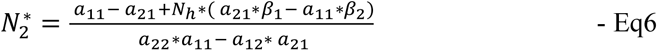

where *N*_1_^∗^ and *N*_2_^∗^ are the population sizes of the two species that reach equilibria in the presence of herbivores. Because intraspecific competition is often higher than interspecific competition (*α_ii_* > *α_ij_*), herbivory will increase or decrease the population growth of the focal species (species 1) depending on the positive or negative values of *α*_12_ ∗ *β*_2_ − *α*_22_ ∗ *β*_1_. For example, herbivory increases the population size of species 1, if *α*_12_ ∗ *β*_2_ − *α*_22_ ∗ *β*_1_ > 0, which can be expressed as 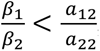, and the two species coexist (1 – *α* _12_ *N*_2_ – *β*_1_ *N*_h_ > 0 & 1 − *α*_21_ *N*_1_ − *β*_2_*N*_ℎ_ > 0). This can be interpreted as when the ratio of herbivory effects on the two species is less than the ratio of competitiveness of species 2 against species 1 and intraspecific competition of 2, and the combined effects of herbivory and interspecific competition are smaller than 1. Similar conditions also apply to species 2.

### Testing model predictions using laboratory microcosms

Feeding our theoretic model with experimental data of model-guided two-species assays allowed us to estimate both intra- and interspecific competition coefficients, as well as individual consumption rates (Fig. 1b). Here, we first grew duckweed and algae either separately or together in beakers. Overall, the interspecific competition (in the absence of herbivory) between algae and duckweed was asymmetric (SI Fig. S3a-b). Although algae strongly reduced duckweed growth, duckweed did not significantly affect algal growth.

Second, quantifying consumption rates (in the absence of a competitor) of snails on duckweed and on algae suggested that snail herbivory reduced the growth of both algae and duckweed (SI Fig. S3c-d); however, the effect of herbivory on algae was greater than that on duckweed. Based on the estimated parameters and coefficients (Fig. 1b), our model predicted that duckweed and algae would coexist in the absence of snails, but the duckweed population would remain small (SI Fig. S4). However, in the presence of a snail, duckweed will perform better and outcompete algae if the starting algal population size is small or intermediate (SI Fig. S4). Our model also predicted that if the number of snails was too high (greater than two) when using our experimental laboratory setup using beakers, both the duckweed and *C. reinhardii* populations would collapse.

To test these predictions, we grew duckweed and algae in the presence and absence of snails for 14 days in indoor three-species microcosms using beakers. As predicted, two snails often extinguished the populations. Therefore, we focused on comparisons between one and no snails. In three out of the four scenarios, where the algae density was relatively low or intermediate high, snails indeed increased the duckweed growth rate (*P≤0.02,* Wilcoxon test, n=3-8, Fig. 1c). However, when the starting algae density was very high (5.8 [10^6^ cells mL^-1^]) and the starting duckweed population size was low (100 fronds), snail herbivory also led to the collapse of the duckweed population (Fig. 1c).

Our theoretic model suggests that *S. polyrhiza* benefits under herbivory in the presence of *C. reinhardii*, when 1 − *α*_12_ ∗ *N*_2_ ∗ *β*_1_ ∗ *N*_ℎ_ > 0. The outcome of the model prediction is consistent with the experimental result in three out of the four scenarios when incorporating the 95 % confidence interval of bootstrapped coefficients (Fig. 1c).

Our model also predicts that snails always reduce algal population size 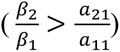 (Fig. 1b). In our experiments, the algal populations grew significantly worse in the presence of snails under nearly all tested conditions (*P≤0.03,* Wilcoxon test, n=3-8, SI Fig. S5).

### Quantifying herbivory effects in outdoor mesocosms

We tested whether snail herbivory can increase duckweed growth via reduced competition under outdoor conditions by growing replicated monoclonal duckweed populations and their native algal community in the presence and absence of snails across one growth season in outdoor ponds (Fig. 2a-b). Three weeks after the start of the experiments, snail-infested duckweed population covered an average of 57 % larger surface than control populations, and this difference even grew towards the end of the experiment (*P*=0.000002, *X*^2^=22.5, linear-mixed effect model, Fig. 2c-d). Increased duckweed coverage occurred despite the duckweeds being attacked by the snails, which was evident from visual damage and the increase in the levels of phenylpropanoids, particularly of flavonoids (SI Table S5). The increased duckweed surface coverage cannot be explained by the increased biomass per frond, as biomass per frond was reduced rather than increased under herbivory at week four (*P*=0.002, linear mixed effect model, SI Fig. S5) with diminishing effects in the following week. Increased duckweed growth under snail herbivory was likely not due to altered nutrient levels, as nutrient levels did not differ significantly between the adjacent snail and herbivore compartments (SI Table S4).

**Figure 2.**
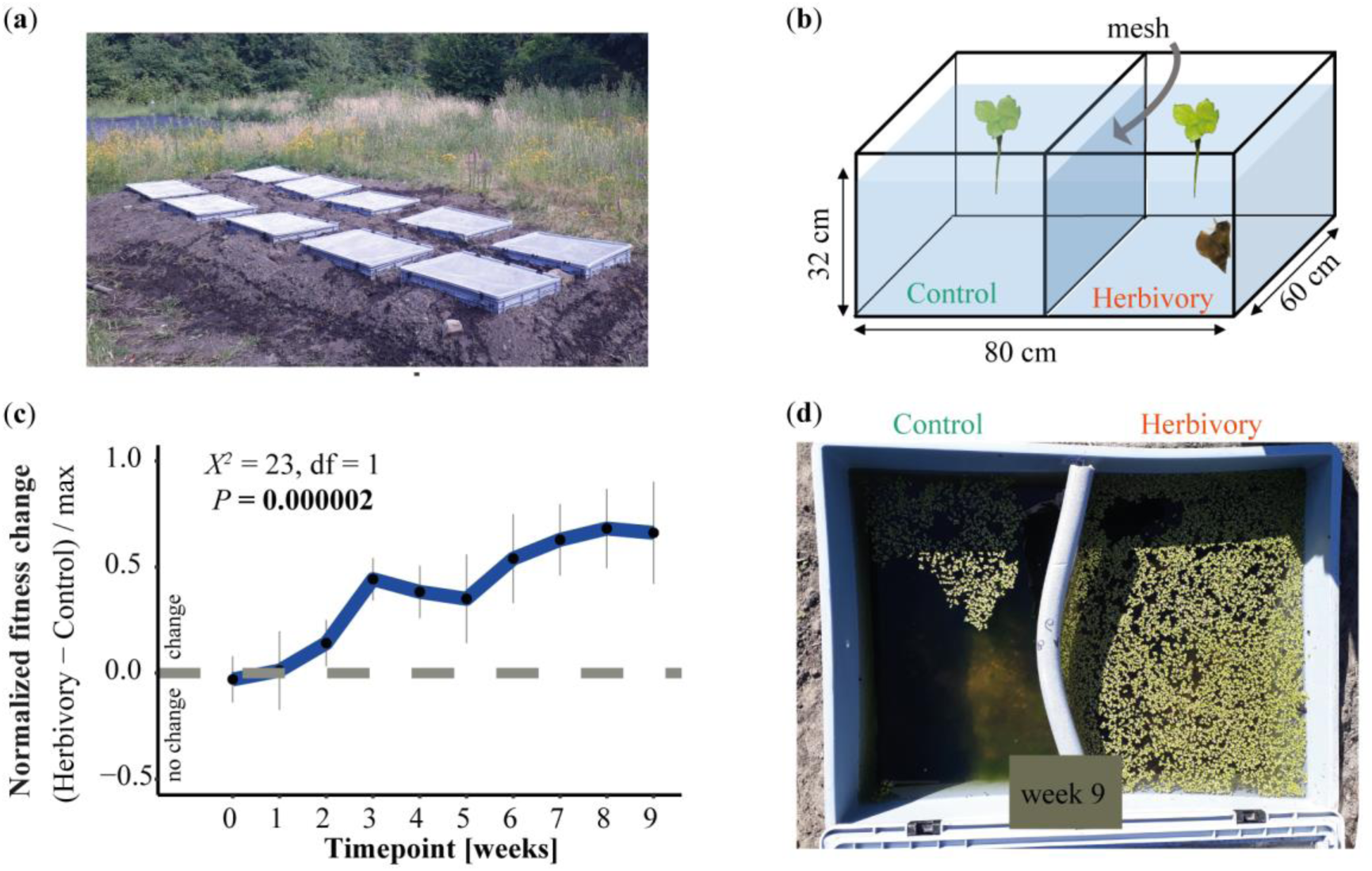
Herbivory treatment increased relative fitness of *Spirodela polyrhiza* in outdoor mesocosms. (**a**) Field site with eight replicated ponds covered with mesh. (**b**) Schematic representation of each pond. Treatment compartments with presence and absence of the snail herbivore *Lymnaea stagnalis* were separated by a mesh, preventing snails to enter the control compartment while allowing for water exchange. (**c**) Presence of *L. stagnalis* increased the fitness of *S. polyrhiza* outdoors over a course of nine weeks during one summer season representing about 15-20 asexual generations of plants (n=4-5, linear mixed effect model). Values above 0 indicate herbivory increased duckweed fitness. Y-axis refers to normalized fitness and x-axis refers to time. (**d**) Representative plant populations in week nine.

Shortly after setting up the experiment, we observed rapid growth of microalgae, concurrent with an increase in water and air temperature (SI Fig. S6), with a peak at week four. Based on morphology, we identified *Chlamydomonas* ssp. as the dominant microalgal species, accompanied by *Closterium* sp. and *Lagynion* sp. (Fig. 3a). Amplicon metagenomics using the 18S rRNA gene marker on duckweed samples with adherent microalgae revealed that the dominant microalgae were Chlamydomonadales (SI Fig. S7). Herbivory treatment did not affect microalgal and cyanobacterial community structure (SI Fig. S8a-b), neither the microbial diversity associated with *S. polyrhiza* fronds (*P*>0.2, PERMANOVA, SI Figure S7, SI Table S6) nor the relative abundance of different algal genera (Fig. 3c-d). However, our method was unable to estimate the absolute quantity of microalgae.

**Figure 3.**
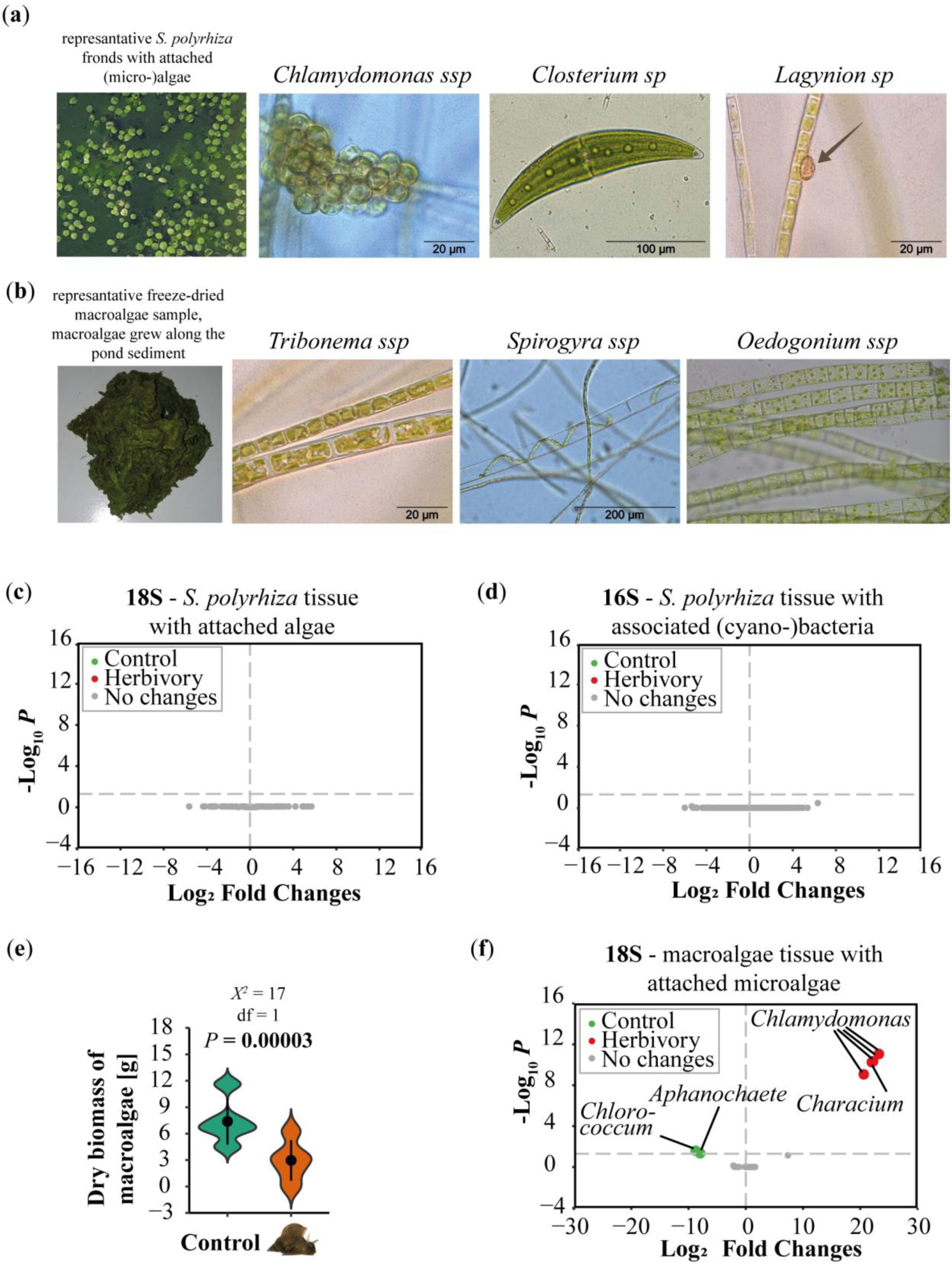
Algae community lived together with *Spirodela polyrhiza* in outdoor mesocosms. Herbivory treatment reduced abundance of macroalgae biomass and partly affected the abundance of microalgae genera associated with macroalgae samples in outdoor ponds. (**a**) In early summer abundance of microalgae was high. Representative pictures of dominant microalgae species, later confirmed via amplicon metagenomic approach. (**b**) During the second half of the season, macroalgae started growing. Representative pictures of dominant macroalgae species harvested at the end of the season, later confirmed via amplicon metagenomic approach. (**c**) Presence of snails did not affect the abundance of different genera of algae (18S) and (**d**) (cyano-)bacteria (16S) associated with *S. polyrhiza* samples. (**e**) Presence of snails significantly reduced macroalgae abundance (linear mixed effect model, n=5). (**f**) Herbivory affected the relative abundance of 18S genus hits associated with macroalgae samples. Higher abundance of genera in either control (left green) or herbivory (right red) treatment. Microscopic pictures were taken by Antje Gutowski.

In the second half of the experimental period (weeks 4–9), thick clumps of filamentous macroalgae started to grow along the bottom of the pond (Fig. 3b) and eventually started to attach to the duckweed roots. At the end of the growing season, these macroalgae were significantly less abundant in the presence than in the absence of the snail herbivore (*P*=0.00003, linear mixed-effect model, Fig. 3e). An additional feeding experiment also showed that the snails could directly feed on the macroalgae collected from the ponds (SI Fig. S9). Based on morphology, we identified *Tribonema* ssp., *Oedogonium* ssp., and *Spirogyra* ssp. (Fig. 3b) as the dominant macroalgal species, which was further confirmed by 18S amplicon sequencing (SI Fig. S10). Again, based on 18S rRNA gene marker, Chlamydomonadales was the dominant microalga associated with macroalgae samples. Shannon diversity of macroalgae with adherent microalgae was significantly higher under herbivory (*P*=0.03, linear mixed effect model, SI Fig. S10), whereas community structure was unaffected by herbivory (SI Fig. S8c). Furthermore, herbivory affected the abundance of different genera associated with macroalgae samples, for example, a higher relative abundance under herbivory for *Chlamydomonas* (Fig. 3f).

Together, these results are consistent with our model prediction that snails increase duckweed fitness by reducing the growth of their competitors, here the macroalgae, under outdoor conditions.

## Discussion

Herbivory and competition are two major factors that can influence plant fitness. Although these two factors are often studied separately, they are intertwined, as herbivores can alter the competitive landscape when feeding on multiple species. Here, we provide parallel evidence that herbivory can increase plant fitness via reduced interspecific competition. First, we developed a theoretic model to understand the fitness consequences of competing (plant) species under herbivory. We found that herbivores can increase or decrease plant fitness depending on the relative ratio of intra- and interspecific competition as well as the relative consumption rates of focal and competing species, respectively. Second, we tested and validated our model predictions using three aquatic species (including microalgae) growing together in an indoor microcosm. Third, the outdoor mesocosms confirmed that herbivory can increase plant fitness by reducing competing species (here macroalgae) under natural conditions.

We developed our model by introducing plant-herbivore interactions into the existing species competition model. For simplicity, we kept the herbivore population size constant, which mimics natural situations when the herbivore life cycle is much longer than the plant life cycle. This is the case in our duckweed-algae-snail system, and it is similar to other aquatic plant-herbivore interactions. For example, the aquatic nymphal stage of mayflies can last for up to two and a half years while feeding on algae, epiphytes, and aquatic plants with various life spans [36]. In other systems, however, herbivores and plants may reproduce at similar rates or even higher rates, such as aphids feeding on mustard plants [37]. Please note here that within the timeframe of our indoor model-guided assays (which are the source data used to feed our theorical model to then estimate model coefficients) adult *L. stagnalis* did not propagate. Future generic models that include the population dynamics of herbivores (e.g. [38]), as well as mesocosm approaches that manipulate and quantify herbivore densities (as indeed snails laid eggs in our outdoor mesocosms over the course of 9 weeks), will provide further insights into the species dynamics of multitrophic communities.

To determine interspecific competition coefficients, we used model-guided assays in which we kept the abundance of the competitor constant over time while varying the starting abundance of the focal species and measuring the focal species’ per capita growth (fitness) (SI Fig. S3), assuming a linear relationship. While this approach made the experiments in the lab feasible, considering non-linearity within our model, as proposed elsewhere, [9,10,39,40] will further improve the realistic characterization of the species dynamics. While the current model analysis, considering linearity and using bootstrapped coefficients, demonstrates an alignment between the theoretical model and experimental findings in three out of four scenarios (Fig. 1c), exploring alternative methodologies holds promise for further refining and enhancing the robustness of the results.

Despite these limitations, we experimentally validated the predicted outcomes of our model by using three interacting species under indoor conditions. Our results showed that snails benefitted duckweed fitness in the presence of low (0.9 [10^6^ cells mL^-1^]) and intermediate-high (4.1 [10^6^ cells mL^-1^]) starting concentrations of the competing algae, whereas snails had neutral effects on duckweed fitness when competing against high (5.8 [10^6^ cells mL^-1^]) algae concentrations (Fig. 1c). In contrast, algae did not benefit from the presence of duckweed under herbivory in the tested scenarios (SI Fig. S5). In nature, snails feed on both micro- and macroalgae, which have also been found in outdoor communities [41–44]. It is likely that snails prefer algae over duckweed, as macroalgae have thinner and more easily palatable cell walls compared to macrophytes [41].

Under outdoor conditions, the algal community changed over time, which also likely led to changes in intra- and interspecific competition between algae and duckweed. Therefore, it is technically challenging to quantify model parameters directly *in situ*. Nevertheless, we found evidence that herbivory can indeed increase duckweed fitness by reducing algal abundance outdoors, which is likely due to the combined effects of the relative changes and absolute reduction of both micro- and macro-algae. Duckweed can suppress algae growth by reducing light availability [19] and nutrients [45], while in turn, algae can produce toxins and increase the pH to levels toxic to duckweeds [19,46]. Although we did not attempt to elucidate which of these mechanisms shaped duckweed-algae competition, our findings are in line with other studies, in which snails increased the growth of floating and submerged plants [47] and reduced the abundance of epiphytic communities thereby benefitting macrophytes [42,48]. Furthermore, it was shown before that the removal of macroalgae (e.g., *Chara* sp. and *Vaucheria* sp.) can increased the growth of a lakés macrophytes [49]. Similar effects were also observed in terrestrial plant-herbivore interactions. For example, introducing novel herbivores to a plant community enhanced the growth of small-stature plants by reducing the dominant plant species and increasing light availability [50]. These studies and our results highlight the importance of considering indirect effects, such as competition, when studying plant-herbivore interactions. Therefore, from an evolutionary perspective, herbivores cannot only drive the evolutionary changes in traits involved in defences but also shape the evolution of traits that are associated with intra- and interspecific competition in nature.

Here, we demonstrate that herbivores determine plant fitness through both direct negative effects through feeding and indirect beneficial effects through the reduced growth of competitors. Consequently, herbivores can increase plant fitness by reducing the abundance of the competing species. Our model allows the quantification of these effects, showing that plant fitness depends on the relative ratio of intra- and interspecific competition, as well as the relative consumption rates of focal and competing species, respectively. Our study highlights that plant-herbivore interactions and their evolution should be studied in a community context, as indirect effects can sometimes overwrite direct effects.

## Data accessibility statement

The datasets and code supporting this article have been uploaded as part of the supplementary material. Raw sequencing data is available on NCBI SRA under BioProject PRJNA914505 (16S duckweed samples), PRJNA914509 (18S duckweeds samples), PRJNA996071 (18S algae samples). The code to replicate sequencing data analyses, as well as raw data for our modelling approach are available at https://github.com/Xu-lab-Evolution/Duckweed-Algae-Snail_Project.

## Acknowledgements

We thank for experimental assistances from Marina Hölter, Mark Winzen, Joachim Rikus, Dennis Zascerinskij, Zahra Kouzbour, Maximilian Schiefer, Tom Lieth, Marie Sárazová, and Ursula Martiné. We thank Antje Gutowski from AlgaLab Bremen for her support to morphologically identify algae species and for providing pictures of our algae samples. We also thank Holger Schön for crafting the outdoor ponds, and Maik Bartelheimer and Ortrun Lepping for providing the initial *Chlamydomonas* culture. We thank Simon Hart for the discussion of our modelling approach. We thank Elisabeth Meyer for sharing the pond equipment. We thank Otto Klemm from the University of Münster for providing weather data.

## References

1. Sala OE et al. 2000 Global biodiversity scenarios for the year 2100. Science 287, 1770–1774. (doi:10.1126/SCIENCE.287.5459.1770)

2. Pereira HM et al. 2010 Scenarios for global biodiversity in the 21st century. Science (1979) 330, 1496–1501. (doi:10.1126/science.1196624)

3. Hooper DU et al. 2012 A global synthesis reveals biodiversity loss as a major driver of ecosystem change. Nature 2012 486:7401 486, 105–108. (doi:10.1038/nature11118)

4. Allan E, Crawley MJ. 2011 Contrasting effects of insect and molluscan herbivores on plant diversity in a long-term field experiment. Ecol Lett 14, 1246–1253. (doi:10.1111/J.1461-0248.2011.01694.X)

5. Carson WP, Root RB. 2000 Herbivory and plant species coexistence: community regulation by an outbreaking phytophagous insect. Ecol Monogr 70, 73–99. (doi:10.1890/0012-9615)

6. Stevens H. 2021 Primer of Ecology using R. See https://hankstevens.github.io/Primer-of-Ecology/comp.html (accessed on 5 February 2023).

7. Lamonica D, Clément B, Charles S, Lopes C. 2016 Modelling algae–duckweed interaction under chemical pressure within a laboratory microcosm. Ecotoxicol Environ Saf 128, 252–265. (doi:10.1016/j.ecoenv.2016.02.008)

8. Lamonica D, Herbach U, Orias F, Clément B, Charles S, Lopes C. 2016 Mechanistic modelling of daphnid-algae dynamics within a laboratory microcosm. Ecol Modell 320, 213–230. (doi:10.1016/J.ECOLMODEL.2015.09.020)

9. Hart SP, Turcotte MM, Levine JM. 2019 Effects of rapid evolution on species coexistence. Proc Natl Acad Sci U S A 116, 2112–2117. (doi:10.1111/nph.18709)

10. Hart SP, Freckleton RP, Levine JM. 2018 How to quantify competitive ability. Journal of Ecology 106, 1902–1909. (doi:10.1111/1365-2745.12954)

11. Hess C, Levine JM, Turcotte MM, Hart SP. 2022 Phenotypic plasticity promotes species coexistence. Nat Ecol Evol 6, 1256–1261. (doi:10.1038/s41559-022-01826-8)

12. Levine JM, Bascompte J, Adler PB, Allesina S. 2017 Beyond pairwise mechanisms of species coexistence in complex communities. Nature. 546, 56–64. (doi:10.1038/nature22898)

13. Erb M. 2018 Plant defenses against herbivory: closing the fitness gap. Trends Plant Sci 23, 187–194. (doi:10.1016/j.tplants.2017.11.005)

14. Agrawal AA, Maron JL. 2022 Long-term impacts of insect herbivores on plant populations and communities. Journal of Ecology 110, 2800–2811. (doi:10.1111/1365-2745.13996)

15. Pardo I, Doak DF, García-González R, Gómez D, García MB. 2015 Long-term response of plant communities to herbivore exclusion at high elevation grasslands. Biodivers Conserv 24, 3033–3047.

16. Petruzzella A, van Leeuwen CHA, van Donk E, Bakker ES. 2020 Direct and indirect effects of native plants and herbivores on biotic resistance to alien aquatic plant invasions. Journal of Ecology 108, 1487–1496. (doi:10.1111/1365-2745.13380)

17. Landolt E. 1957 Physiologische und ökologische Untersuchungen an Lemnaceen. Berichte der Schweizerischen Botanischen Gesellschaft 67, 271–410.

18. Ziegler P, Adelmann K, Zimmer S, Schmidt C, Appenroth KJ. 2015 Relative *in vitro* growth rates of duckweeds (Lemnaceae) - the most rapidly growing higher plants. Plant Biol 17, 33–41. (doi:10.1111/plb.12184)

19. Roijackers R, Szabó S, Scheffer M. 2004 Experimental analysis of the competition between algae and duckweed. Arch Hydrobiol 160, 401–412. (doi:10.1127/0003-9136/2004/0160-0401)

20. Edwards P, Hassan MS, Chao CH, Pacharaprakiti C. 1992 Cultivation of duckweeds in septage-loaded earthen ponds. Bioresour Technol 40, 109–117. (doi:10.1016/0960-8524(92)90195-4)

21. Bernd KK, Cook N. 2002 Microscale assay monitors algal growth characteristics. Biotechniques 32, 1256–1259. (doi:10.2144/02326BM08)

22. Hart SP, Turcotte MM, Levine JM. 2019 Effects of rapid evolution on species coexistence. Proc Natl Acad Sci U S A 116, 2112–2117. (doi:10.1073/pnas.1816298116)

23. Fodor I, Hussein AAA, Benjamin PR, Koene JM, Pirger Z. 2020 The unlimited potential of the great pond snail, *Lymnaea stagnalis*. Elife 9, 1–18. (doi:10.7554/ELIFE.56962)

24. Appenroth KJ, Teller S, Horn M. 1996 Photophysiology of turion formation and germination in *Spirodela polyrhiza*. Biol Plant 38, 95–106. (doi:10.1007/BF02879642)

25. R Core Team. 2022 R: A Language and Environment for Statistical Computing.

26. Wickham H. 2016 ggplot2: Elegant Graphics for Data Analysis.

27. Schneider CA, Rasband WS, Eliceiri KW. 2012 NIH Image to ImageJ: 25 years of image analysis. Nature Methods 2012 9:7 9, 671–675. (doi:10.1038/nmeth.2089)

28. Callahan BJ, McMurdie PJ, Rosen MJ, Han AW, Johnson AJA, Holmes SP. 2016 DADA2: High-resolution sample inference from Illumina amplicon data. Nature Methods 2016 13:7 13, 581–583. (doi:10.1038/nmeth.3869)

29. Quast C, Pruesse E, Yilmaz P, Gerken J, Schweer T, Yarza P, Peplies J, Glöckner FO. 2013 The SILVA ribosomal RNA gene database project: improved data processing and web-based tools. Nucleic Acids Res 41, D590–D596. (doi:10.1093/NAR/GKS1219)

30. Guillou L et al. 2013 The Protist Ribosomal Reference database (PR2): a catalog of unicellular eukaryote small sub-unit rRNA sequences with curated taxonomy. Nucleic Acids Res 41. (doi:10.1093/NAR/GKS1160)

31. McMurdie PJ, Holmes S. 2013 phyloseq: An R Package for Reproducible Interactive Analysis and Graphics of Microbiome Census Data. PLoS One 8, e61217. (doi:10.1371/JOURNAL.PONE.0061217)

32. Dixon P. 2003 VEGAN, a package of R functions for community ecology. Journal of Vegetation Science 14, 927–930. (doi:10.1111/J.1654-1103.2003.TB02228.X)

33. Love MI, Huber W, Anders S. 2014 Moderated estimation of fold change and dispersion for RNA-seq data with DESeq2. Genome Biol 15, 1–21. (doi:10.1186/S13059-014-0550-8/FIGURES/9)

34. Bates D, Mächler M, Bolker B, Walker S. 2015 Fitting Linear Mixed-Effects Models Using lme4. Jouranal of Statistical Software 67, 1–48.

35. Davis NM, Proctor DiM, Holmes SP, Relman DA, Callahan BJ. 2018 Simple statistical identification and removal of contaminant sequences in marker-gene and metagenomics data. Microbiome 6, 1–14. (doi:10.1186/S40168-018-0605-2/FIGURES/6)

36. Brittain JE. 1982 Biology of mayflies. https://doi.org/10.1146/annurev.en.27.010182.001003 27, 119–147. (doi:10.1146/ANNUREV.EN.27.010182.001003)

37. Gautam MP, Patel SV, Singh SN, Kumar P, Yadav SK, Singh DP, Pandey MK. 2019 Mustard aphid, Lipaphis erysimi (Kalt) (Hemiptera: Aphididae): A review. The Pharma Innovation Journal 8, 90–95.

38. Chesson P, Kuang JJ. 2008 The interaction between predation and competition. Nature 456, 235–238. (doi:10.1038/nature07248)

39. Spaak JW, Millet R, Ke PJ, Letten AD, De Laender F. 2023 The effect of non-linear competitive interactions on quantifying niche and fitness differences. Theor Ecol 16, 161–170. (doi:10.1007/S12080-023-00560-6/FIGURES/3)

40. Hart SP, Usinowicz J, Levine JM. 2017 The spatial scales of species coexistence. Nature Ecology & Evolution 2017 1:8 1, 1066–1073. (doi:10.1038/s41559-017-0230-7)

41. Rybak AS. 2016 Freshwater population of Ulva flexuosa (Ulvaceae, Chlorophyta) as a food source for great pond snail: *Lymnaea stagnalis* (Mollusca, Lymnaeidae). Phycological Res 64, 207–211. (doi:10.1111/PRE.12138)

42. Brönmark C. 1989 Interactions between epiphytes, macrophytes and freshwater snails: A review. Journal of Molluscan Studies 55, 299–311. (doi:10.1093/mollus/55.2.299)

43. Elger A, Lemoine D. 2005 Determinants of macrophyte palatability to the pond snail *Lymnaea stagnalis*. Freshw Biol 50, 86–95. (doi:10.1111/j.1365-2427.2004.01308.x)

44. Brönmark C, Rundle SD, Erlandsson A. 1991 Interactions between freshwater snails and tadpoles: competition and facilitation. Oecologia 87, 8–18. (doi:10.1007/BF00323774)

45. Szabó S, Roijackers R, Scheffer M, Borics G. 2005 The strength of limiting factors for duckweed during algal competition. Arch Hydrobiol 164, 127–140. (doi:10.1127/0003-9136/2005/0164-0127)

46. Landolt E. 1986 Biosystematic Investigations in the Family of Duckweeds (Lemnaceae). The Family of Lemnaceae. A Monographic Study.

47. Koleszár G, Nagy Z, Peeters ETHM, Borics G, Várbíró G, Birk S, Szabó S. 2022 The role of epiphytic algae and grazing snails in stable states of submerged and of free-floating plants. Ecosystems 25, 1371–1383. (doi:10.1007/s10021-021-00721-w)

48. Mormul RP, Ahlgren J, Brönmark C. 2018 Snails have stronger indirect positive effects on submerged macrophyte growth attributes than zooplankton. Hydrobiologia 807, 165–173. (doi:10.1007/s10750-017-3391-0)

49. Vejříková I, Vejřík L, Lepš J, Kočvara L, Sajdlová Z, Čtvrtlíková M, Peterka J. 2018 Impact of herbivory and competition on lake ecosystem structure: underwater experimental manipulation. Scientific Reports 2018 8:1 8, 1–10. (doi:10.1038/s41598-018-30598-0)

50. Descombes P, Pitteloud C, Glauser G, Defossez E, Kergunteuil A, Allard P-M, Rasmann S, Pellissier L. 2020 Novel trophic interactions under climate change promote alpine plant coexistence. Science (1979) **370**, 1469–1473. (doi:10.1126/science.abd7015)

