## Supplemental for "Herbivory can increase plant fitness via reduced interspecific competition – evidence from models and mesocosms"

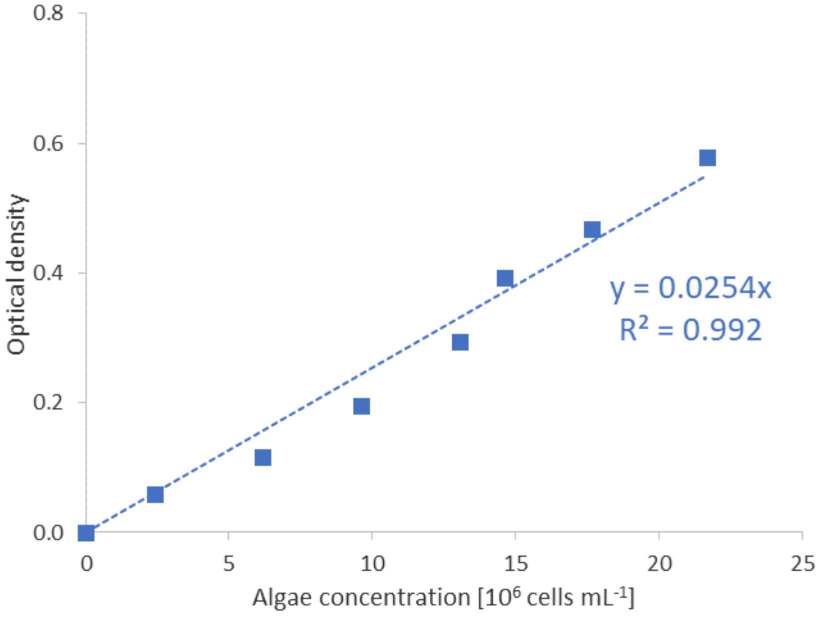


**Figure S1 Reference curve showing the relationship between optical density (OD) and cell concentration of Chlamydomonas reinhardii in Mix-Medium at wavelength 750 nm.** Cell numbers were based on counts using a Thoma chamber. Linear regression forced through the origin was used.

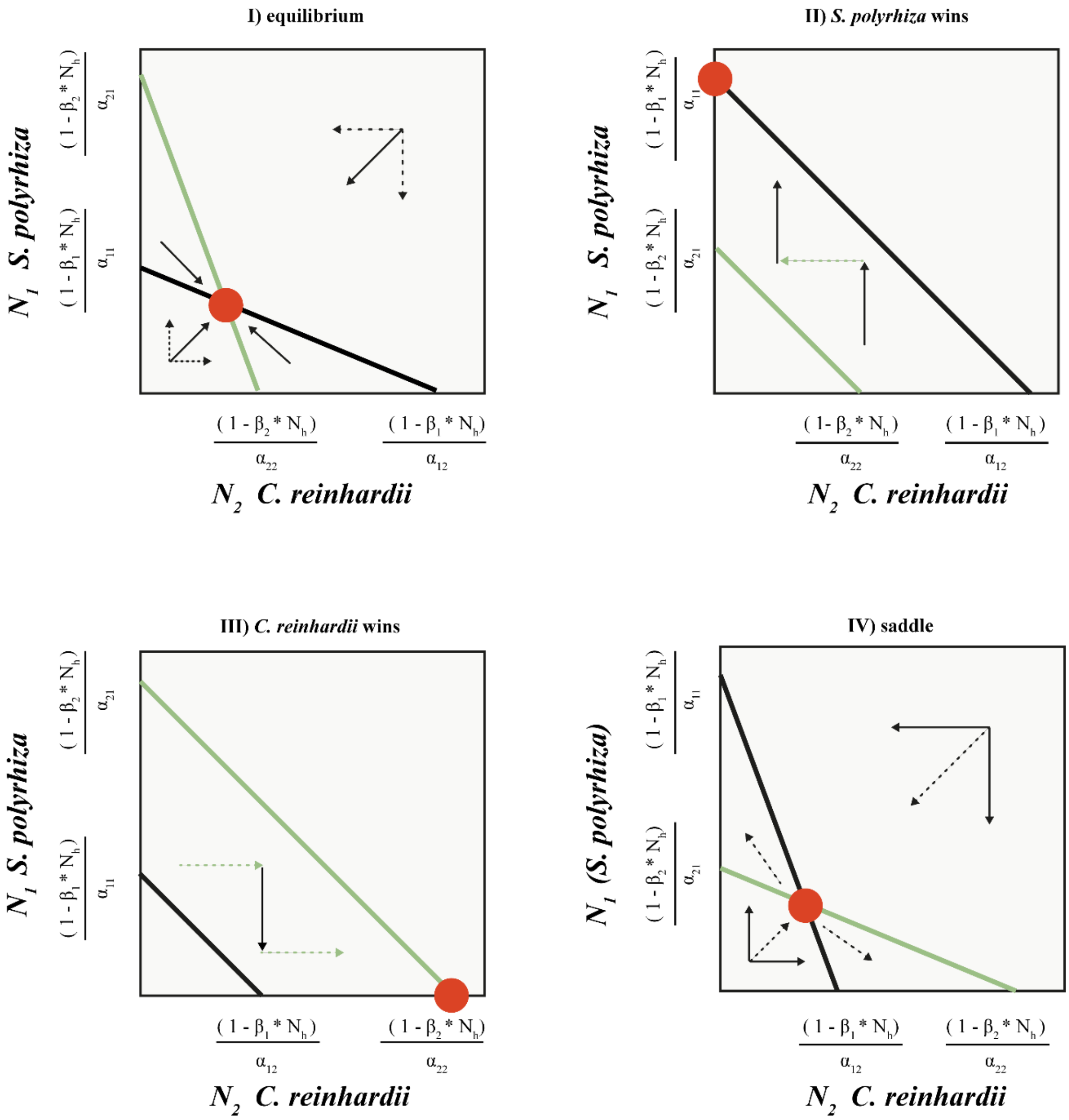


**Figure S2 Phase plane diagrams of four theoretic scenarios describing the interaction of *S. polyrhiza* and *C. reinhardii* under herbivor**y. Horizontal and vertical arrows indicate directions of attraction and repulsion for each population (solid and dashed arrows); diagonal arrows indicate combined trajectory. Circles indicate equilibria; additional boundary equilibria can occur whenever one species is zero. Diagrams are adjusted from (Stevens, 2021).

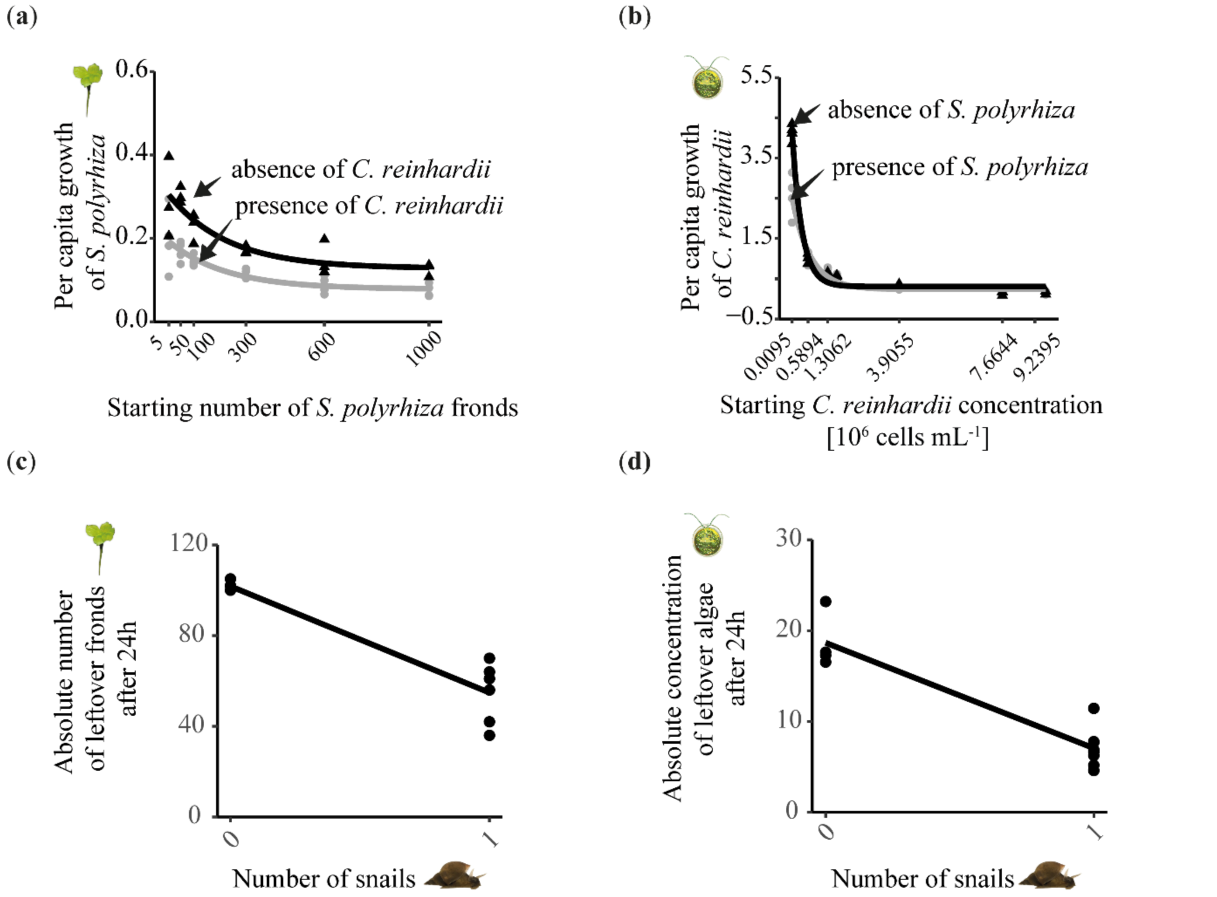


**Figure S3 Model-guided reductionistic 2-species microcosm experimental data used to estimate coefficients of theoretic model.** (**a-b**) Per capita growth measured within 7 days, stat_smooth(method = "nls", formula = y ~ SSasymp(x, Asym, R0, lrc), se = FALSE), (**c-d**) Absolute consumption rates measured within 24 h, geom_smooth(method = lm). (**c**) Starting number of fronds was always 100. (**d**) Starting concentration of algae was always 12.6 [$10^6$ cells $mL^{-1}$]. For details see also the SI Markdown script.

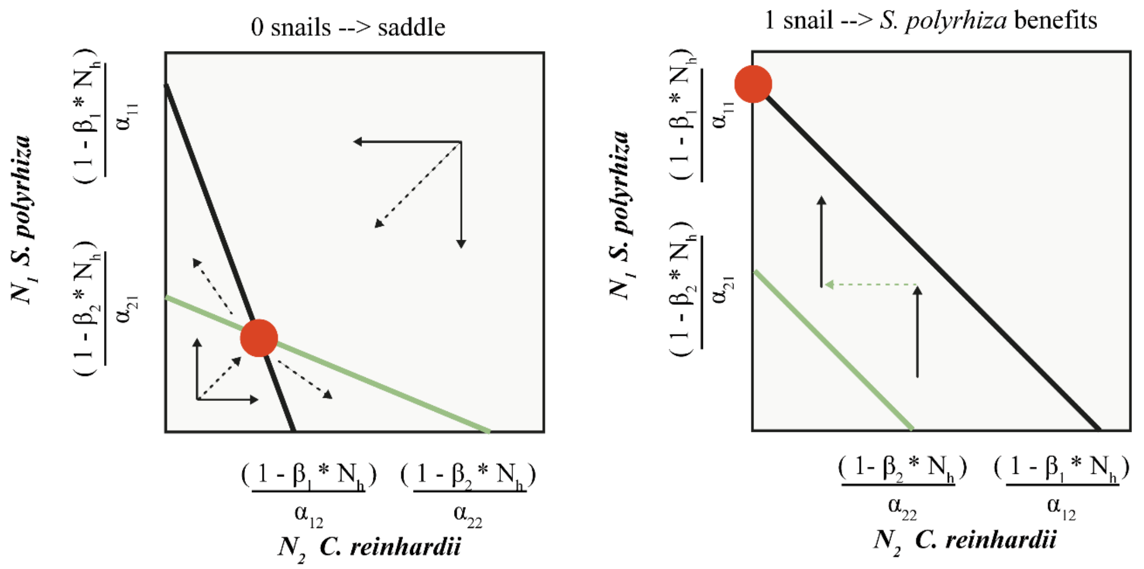


**Figure S4 Scenarios holding true when using our estimated model coefficients.** Our model predicted that duckweed and algae would coexist in the absence of snails, but the duckweed population would remain small (left panel). However, in the presence of a snail, duckweed will perform better and outcompete algae if the starting algal population size is small or intermediate (right panel). Horizontal and vertical arrows indicate directions of attraction and repulsion for each population (solid and dashed arrows); diagonal arrows indicate combined trajectory. Circles indicate equilibria; additional boundary equilibria can occur whenever one species is zero. Diagrams are adjusted from (Stevens, 2021).

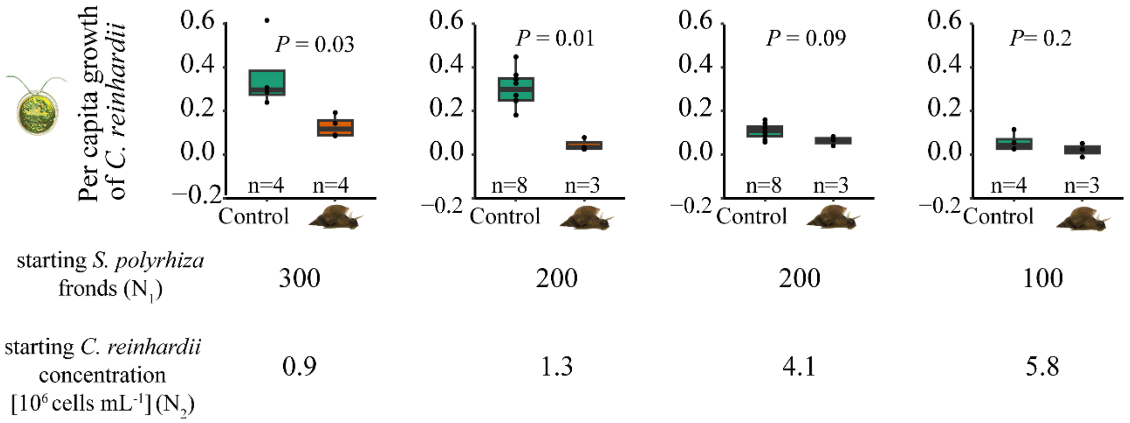


**Figure S5 Indoor microcosm data of algae**. Here, regardless of the starting abundances of the competing species, algae never benefitted from the presence of herbivory when competing with *S. polyrhiza*. *P*-values refer to Wilcoxon tests. Compare with main Figure 1c for *S. polyrhiza* fitness.

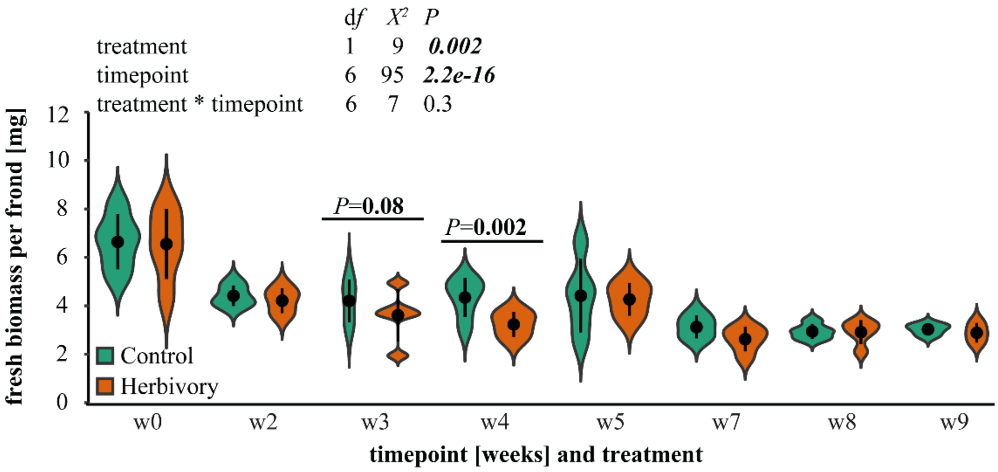


**Figure S5 Biomass data of *S. polyrhiza* fronds over time when grown outdoors in mesocosms.** Pairwise comparisons of fresh weight per frond [mg] between control and snail herbivory treatment over a course of nine weeks outdoors. Biomass per frond was significantly reduced in week 4 but differences diminished later on. *P*-values correspond to linear-mixed effect model with time (for overall model displayed on top) and pond as random factor. Only *P*-values <0.1 are shown when comparing treatments within one timepoint, n=5 except for week 9 n=4.

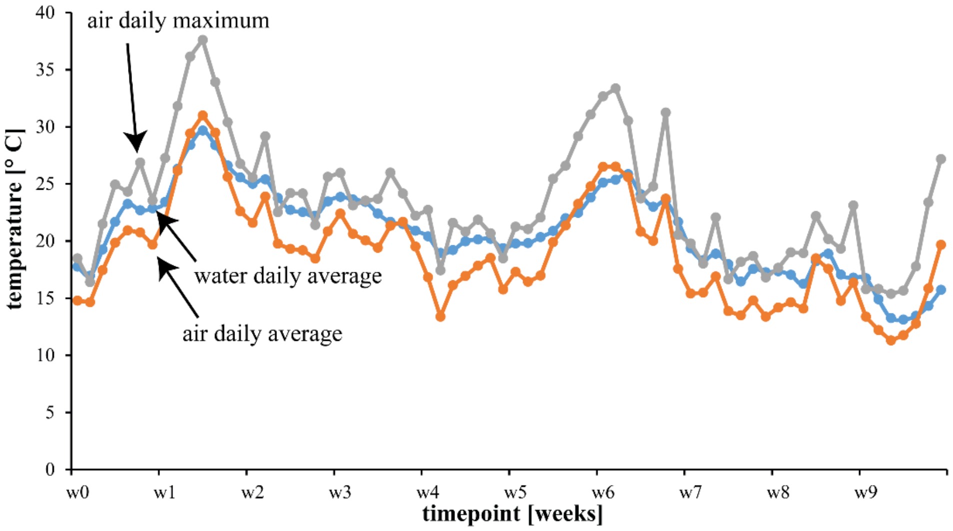


**Figure S6 Daily average water (blue), daily average air (orange) and daily maximum air (grey) temperature of outdoor mesocosm ponds in 2019**. Water temperature was measured with a data logger placed in one of the ponds (HOBO MX2202, Onset Computer Corporation, Bourne, MA, USA). Air temperature data was provided by a nearby weather station.

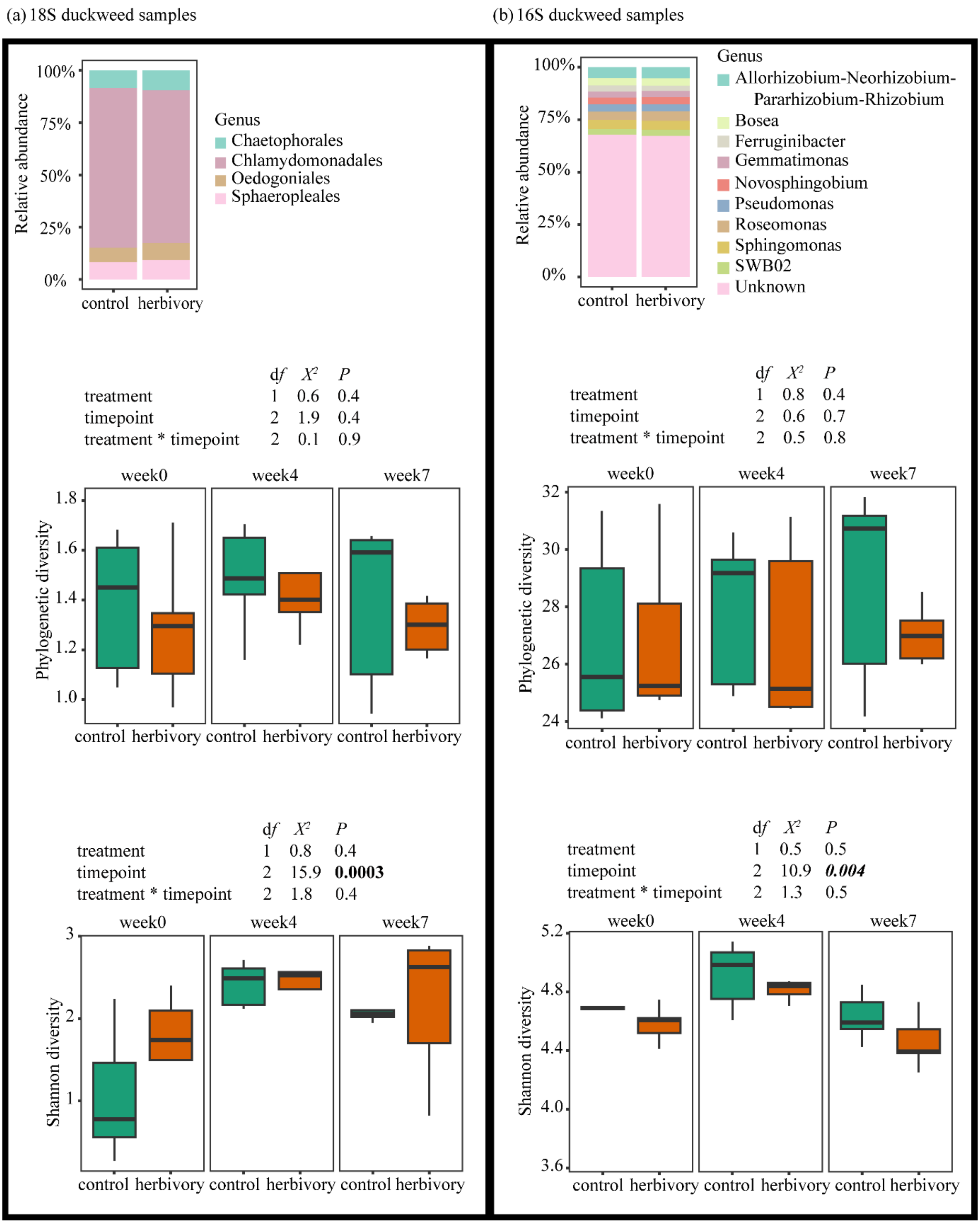
(a) 18S duckweed samples
(b) 16S duckweed samples
Relative abundance
100%
75%
50%
25%
0%
control herbivory
Genus
Chaetophorales
Chlamydomonadales
Oedogoniales
Sphaeropleales
Genus
Allorhizobium-Neorhizobium-Pararhizobium-Rhizobium
Bosea
Ferruginibacter
Gemmatimonas
Novosphingobium
Pseudomonas
Roseomonas
Sphingomonas
SWB02
Unknown
df X² P
treatment 1 0.6 0.4
timepoint 2 1.9 0.4
treatment * timepoint 2 0.1 0.9
df X² P
treatment 1 0.8 0.4
timepoint 2 0.6 0.7
treatment * timepoint 2 0.5 0.8
week0
week4
week7
Phylogenetic diversity
1.0
1.2
1.4
1.6
1.8
24
26
28
30
32
control herbivory control herbivory control herbivory
df X² P
treatment 1 0.8 0.4
timepoint 2 15.9 0.0003
treatment * timepoint 2 1.8 0.4
df X² P
treatment 1 0.5 0.5
timepoint 2 10.9 0.004
treatment * timepoint 2 1.3 0.5
Shannon diversity
1
2
3
3.6
4.0
4.4
4.8
5.2

**Figure S7 Relative abundance and diversity of (a) Algal (18S) and (b) bacterial (including cyanobacteria which represented less than 1 % of hits) (16S) microbiome associated with *S. polyrhiza* grown outdoors**. Timepoint but not herbivory treatment affected Shannon diversity of both algal and bacterial microbiome. *P*-values refer to a linear mixed effect model, n=5.

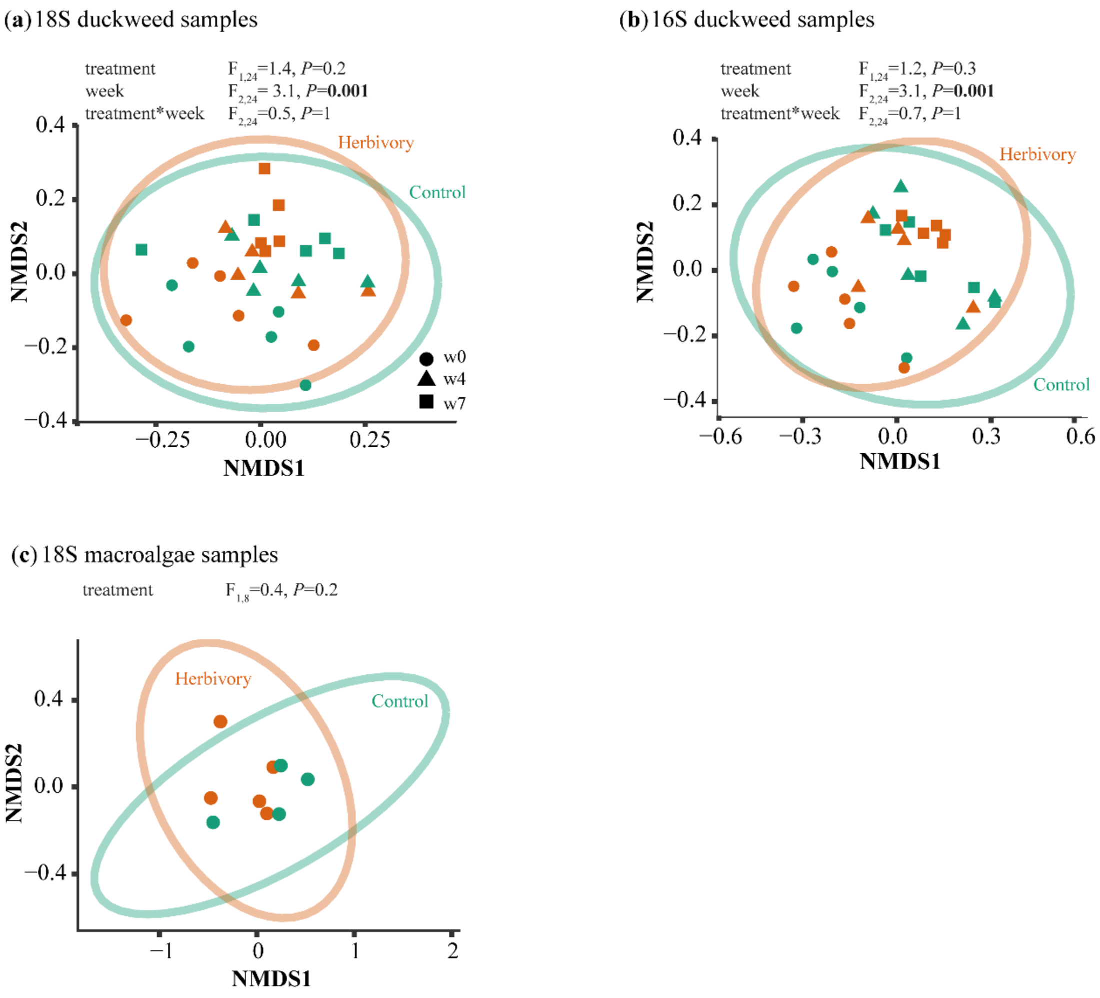


**Figure S8 Community structure of *S. polyrhiza* and macroalgae samples.** (a) Algal community (18S) and (b) (cyano-)bacterial (16S) community structure associated with duckweed samples was not affected by herbivory, but along the season (PERMANOVA, n=5 per timepoint and treatment). (c) Herbivory did not affect community structure of macroalgae samples (including attached microalgae) based on 18S (PERMANOVA, n=4-5).

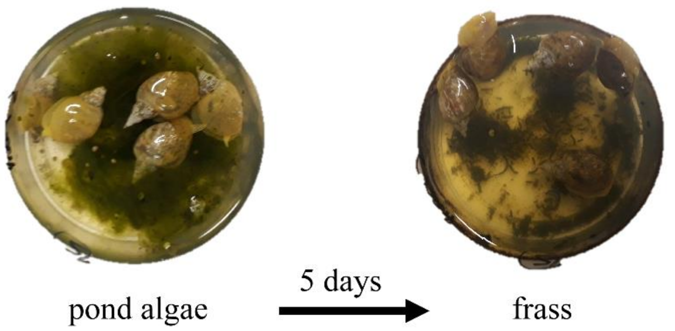


**Figure S9 Representative pictures testing the overall palatability of outdoor harvested macroalgae samples by adult individuals of our herbivorous snail *Lymnaea stagnalis*.** After five days of feeding, only frass was visible (right picture).

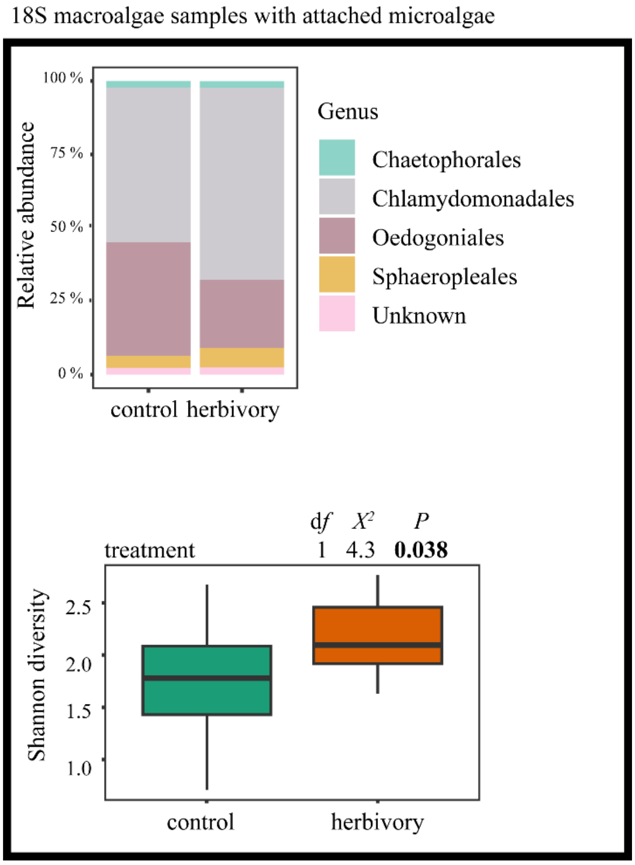


**Figure S10 Relative abundance and diversity of algal (18S) microbiome hits associated with macroalgae samples (including attached microalgae) growing outdoors**. *P*-values refer to a linear mixed effect model, n=4-5.

**Table S1 Recipe for TAP Medium for cultivation of the green microalgae *Chlamydomonas reinhardii*. Recipe was** based on [2]

| | per 1 l * |
|---|---|
| **Stock solution I** | 25 ml |
| $NH_4Cl$ 1.5 g | |
| $MgSO_4$ x 7 $H_20$ 0.4 g in 100 ml | |
| $CaCl_2$ x 2 $H_20$ 0.2 g | |
| **Stock solution II** | 750 µl |
| K-phosphate buffer 0.5 M pH7 | |
| about 30 ml of 0.5 M $KH_2PO_4$ | |
| about 60 ml of 0.5 M $K_2HPO_4$ | |
| **Other** | |
| Acetic acid (100 %, p.a.) | 1 ml |
| Hutner’s trace element solution [3] | 1 ml |
| Tris (solid) | 2.42 g |

*Sterilized and store at 6 °C

**Table S2 Recipe for "Mix-Medium" allowing to cultivate both *Spirodela polyrhiza* and *Chlamydomonas reinhardii* in laboratory microcosms together.** Mixture was based on preliminary experiments allowing survival and growth of both species and acceptance of snail herbivores. Also, N levels of Mix-Medium was chosen to reflect levels of *S. polyrhiza* habitats in nature while increased to10x to ensure growth in a sufficient time-manner.

| | per 1 l | Reference |
|---|---|---|
| **TAP Medium** | 53 ml | Table S1, based on [2] |
| **full N-Medium** | 37 ml | [4] |

**Table S3 Sites from where water was collected for initial inoculation of water body of our outdoor experiment**. Here, we used 1 l per listed location per pond. Water body was prefiltered using a commercial coffee filter, allowing natural microbial communities to enter our pond system.

| Habitat characterization | Location | Coordinates | Occuring genus |
|---|---|---|---|
| Oligotrophic Steam | Haltern am See | 51°44'30.6"N 7°12'17.5"E | *Spirodela, Lemna* |
| Eutrophic pond | Roxel | 51°58'38.4"N 7°29'03.9"E | *Spirodela, Lemna* |
| Mesotrophic pond | Münster | 51°57'40.7"N 7°36'55.7"E | *Spirodela* |

**Table S4 Statistics of nutrient levels in outdoor ponds.** Linear-mixed effects models testing the influence of herbivory on water nutrient levels in outdoor ponds. Data was collected at weeks 0, 3, 5, 7, and 9. Both the pond ID and the sampling timepoints were used as random factors (lmer(Cl ~ treatment * (1|pond) * (1|week)). For multiple comparisons we used Benjamini-Hochberg (BH) corrected p-values. Additionally, PERMANOVA analysis showed no significant differences between herbivory and control treatments ($F_{1,36}$=0.05, P =0.9; 999 permutations stratified over “pond” and “week”), n=4-5.

| **Nutrient** | **df** | $\chi^2$ | ***P*-value** | ***BH corrected P-value*** |
|---|---|---|---|---|
| **Cl** | **1** | 1.6 | 0.2 | 0.5 |
| **$NO_2$** | **1** | 0.2 | 0.6 | 0.1 |
| **$NO_3$** | **1** | 4.1 | 0.04 | 0.1 |
| **$PO_4$** | **1** | 0.3 | 0.6 | 1 |
| **$SO_4$** | **1** | 0.5 | 0.5 | 0.8 |
| **Na** | **1** | 4.5 | 0.03 | 0.5 |
| **$NH_4$** | **1** | 1.2 | 0.3 | 0.8 |
| **K** | **1** | 2.3 | 0.1 | 0.5 |
| **Ca** | **1** | 3.3 | 0.07 | 0.5 |
| **Mg** | **1** | 1.0 | 0.3 | 0.5 |

**Table S5 Statistics of data from metabolite analysis of *S. polyrhiza* fronds cultivated in outdoor mesocosms in 2019**. Samples were collected from control and herbivory compartments in week 0, 3 and 6. [%] refers to percentage of change under herbivory compared to control. Data was analysed using a linear mixed effect model with pond as random factor (lmer(Glycine~treat * (1|Pond), exp2[exp2$week=="w6",]). P-values were further adjusted with Benjamini-Hochberg (p.adjust(p, "BH")), to account for multiple comparisons. N=5, if not stated otherwise.

| # | Metabolite | week0 (n=5) | | | | week3 (n=5) | | | | week6 (n=3) | | | |
|---|---|---|---|---|---|---|---|---|---|---|---|---|---|
| | | % | $\chi^2$ | *P*-value | BH corrected *P*-value | % | $\chi^2$ | *P*-value | BH corrected *P*-value | % | $\chi^2$ | *P*-value | BH corrected *P*-value |
| 1 | L-Alanine | 3 | 0.001 | 1 | 0.7 | 17 | 0.98 | 0.3 | 0.7 | 19 | 1.8678 | 0.2 | 0.3 |
| 2 | L-Arginine | -9 | 1.01 | 0.3 | 1 | 10 | 0.007 | 0.9 | 1 | -23 | 7.67 | **0.0056** | **0.0171** |
| 3 | L-Asparagine | 1 | 3e-04 | 1 | 1 | 2 | 0.003 | 1 | 1 | -27 | 2.91 | 0.09 | 0.2 |
| 4 | L-Aspartic_Acid | 2 | 0.052 | 0.8 | 0.9 | -1 | 0.17 | 0.7 | 0.9 | -5 | 1.988 | 0.2 | 0.3 |
| 5 | L-Glutamic_acid | 8 | 0.24 | 0.6 | 0.9 | 13 | 0.11 | 0.8 | 0.9 | -4 | 0.36 | 0.6 | 0.7 |
| 6 | L-Glutamine | 7 | 0.09 | 0.8 | 0.7 | 20 | 0.85 | 0.4 | 0.7 | -18 | 7.74 | **0.0054** | **0.0171** |
| 7 | L-Isoleucine_1 | 4 | 0.14 | 0.7 | 0.5 | -19 | 2.8 | 0.09 | 0.5 | 36 | 6.9 | **0.008** | **0.0231** |
| 8 | L-Leucine_2 | 1 | 0.017 | 0.9 | 0.4 | -21 | 3.9 | 0.05 | 0.4 | 54 | 3.67 | 0.055 | 0.1 |
| 9 | L-Phenylalanine | 2 | 0.11 | 0.7 | 0.5 | -12 | 2.28 | 0.1 | 0.5 | 12 | 3.8 | 0.051 | 0.1 |
| 10 | L-Proline | -4 | 1.03 | 0.3 | 0.4 | -15 | 4.2 | 0.04 | 0.4 | 11 | 1.22 | 0.3 | 0.4 |

| | | | | | | | | | | | | | |
|---|---|---|---|---|---|---|---|---|---|---|---|---|---|
| 11 | L-Serine | 3 | 0.047 | 0.8 | 0.9 | 11 | 0.14 | 0.7 | 0.9 | 17 | 252.6 | **2.2e-16** | **1.91E-15** |
| 12 | L-Threonine | 4 | 0.4 | 0.5 | 0.3 | -10 | 4.84 | 0.02 | 0.3 | 16 | 5.54 | **0.02** | **0.052** |
| 13 | L-Tryptophane | -3 | 0.62 | 0.4 | 0.7 | -3 | 0.97 | 0.3 | 0.7 | 20 | 2.8 | 0.1 | 0.2 |
| 14 | L-Valine | 0 | 0.038 | 0.9 | 0.7 | -7 | 2.13 | 0.2 | 0.7 | 23 | 3.6 | 0.059 | 0.1 |
| 15 | Ornithine | -6 | 0.9 | 0.3 | 1 | 3 | 0.03 | 0.9 | 1 | -15 | 55.5 | **9.52e-14** | **7.07e-13** |
| 16 | Glycine | -8 | 0.6 | 0.4 | 0.7 | -4 | 0.7 | 0.4 | 0.7 | 1 | 0.005 | 0.9 | 1 |
| 17 | L-Histidine | -4 | 0.76 | 0.4 | 0.8 | 0 | 0.41 | 0.5 | 0.85 | 0 | 7e-04 | 1 | 1 |
| 18 | L-Methionine | -14 | 1.47 | 0.2 | 0.7 | 59 | 0.98 | 0.3 | 0.7 | NA | NA | NA | NA |
| 19 | L-Tyrosine | 3 | 0.03 | 0.9 | 0.4 | -12 | 3.89 | 0.05 | 0.4 | 17 | 1.2 | 0.3 | 0.4 |
| 20 | Tyramine | 28 | 0.29 | 0.6 | 0.8 | 6 | 0.47 | 0.5 | 0.8 | 1 | 1 | 0.3 | 0.4 |
| 21 | Tryptamine | 10 | 1.75 | 0.2 | 0.7 | -19 | 1.61 | 0.2 | 0.7 | 10 | 4 | **0.046** | 0.1 |
| 22 | Shikimic_acid | 10 | 0.41 | 0.5 | 0.9 | 6 | 0.19 | 0.7 | 0.9 | 2 | 0.04 | 1 | 1 |
| 23 | Apigenine | -6 | 0 | 1 | 0.9 | 78 | 0.24 | 0.6 | 0.9 | 46 | 1.4 | 0.2 | 0.3 |
| 24 | Luteoline | 0 | 0.25 | 0.6 | 0.8 | -1 | 0.52 | 0.5 | 0.8 | 42 | 3.1 | 0.07 | 0.1 |
| 25 | IP | -2 | 0.38 | 0.5 | 0.8 | -4 | 0.51 | 0.5 | 0.8 | 18 | 12.9 | **0.0003** | **0.0014** |
| 26 | IPR | 1 | 0.03 | 0.9 | 0.9 | 40 | 0.15 | 0.7 | 0.9 | 6 | 0.3 | 0.6 | 0.7 |
| 27 | tZ | -10 | 6.45 | 0.01 | 0.2 | 32 | 3.37 | 0.01 | 0.2 | 9 | 1.07 | 0.3 | 0.4 |
| 28 | tZR | -5 | 0.59 | 0.44 | 0.7 | 34 | 1.48 | 0.2 | 0.7 | -4 | 0.09 | 0.76 | 0.9 |
| 29 | cZ | -3 | 0.18 | 0.67 | 0.9 | 16 | 0.25 | 0.6 | 0.9 | 15 | 0.002 | 0.97 | 1 |
| 30 | cZR | 0 | 0.06 | 0.8 | 0.9 | -2 | 0.09 | 0.8 | 0.9 | 23 | 0.67 | 0.41 | 0.5 |
| 31 | IAA 130 | 19 | 3.98 | 0.046 | 0.1 | -13 | 9.5 | 0.002 | 0.1 | 26 | 1.98 | 0.16 | 0.3 |

| | | | | | | | | | | | | | |
|---|---|---|---|---|---|---|---|---|---|---|---|---|---|
| 32 | Coumaric_acid_p | 18 | 2.7 | 0.1 | 0.5 | -12 | 2.24 | 0.1 | 0.5 | 16 | 2.48 | 0.11 | 0.2 |
| 33 | Caffeic_acid | 10 | 0.92 | 0.34 | 0.9 | -3 | 0.26 | 0.6 | 0.9 | 13 | 1.85 | 0.17 | 03 |
| 34 | Ferulic_acid | 76 | 0.57 | 0.41 | 0.7 | NA | NA | NA | NA | -7 | 5.77 | 0.02 | **0.052** |
| 35 | Sinapic_acid | 7 | 0.03 | 0.86 | 0.1 | -28 | 8.4 | 0.00 | 0.1 | 0 | 8e-04 | 0.98 | 1 |
| 36 | ABA | -15 | 4.9 | 0.03 | 0.65 | 32 | 1.55 | 0.2 | 0.7 | 0 | 0.01 | 0.9 | 1 |
| 37 | SA | 12 | 1.00 | 0.32 | 0.9 | -3 | 0.23 | 0.63 | 0.9 | 13 | 9.28 | **0.002** | **0.008** |
| 38 | JA | 48 | 0.02 | 0.87 | 1 | 52 | 0.003 | 1 | 1 | 75 | 3.38 | 0.07 | 0.14 |
| 39 | JA-Ile | NA | NA | NA | 1 | 73 | 0.003 | 1 | 1 | 190 | 7.85 | **0.005** | **0.0171** |
| 40 | OPDA | 24* | 0.08* | 0.78* | 1* | 38 | 0.007 | 0.9 | 1 | 92 | 1.89 | 0.17 | 0.3 |
| 41 | Cyanidine_3-Glc | -7 | 1.3 | 0.26 | 0.5 | 23 | 2.3 | 0.1 | 0.5 | 26 | 162.8 | **< 2.2e-16** | **1.91e-15** |
| 42 | Cyanidin-Mal-Glc | -4 | 0.67 | 0.41 | 0.7 | 24 | 1.16 | 0.3 | 0.7 | 26 | 26 | **3.062e-07** | **1.99e-06** |
| 43 | Chlorogenic_acid | 7 | 0.06 | 0.8 | 0.9 | 25 | 0.06 | 0.8 | 0.9 | 52 | 17 | **3.404e-05** | **0.0001** |
| 44 | Luteolin_7-O-Glc | -4 | 0.96 | 0.32 | 0.7 | 8 | 0.83 | 0.4 | 0.7 | 18 | 14 | **0.0001** | **0.0007** |
| 45 | Luteolin_8-C-Glc | -4 | 1 | 0.3 | 0.7 | 10 | 0.9 | 0.3 | 0.7 | 22 | 78 | **2.2e-16** | **1.91e-15** |
| 46 | Apigenin_7-O-Glc | -5 | 0.78 | 0.38 | 0.7 | 12 | 0.76 | 0.4 | 0.7 | 27 | 98 | **2.2e-16** | **1.91e-15** |
| 47 | Apigenin_8-C-Glc | -3 | 0.46 | 0.5 | 0.7 | 12 | 0.73 | 0.4 | 0.7 | 25 | 136 | **2.2e-16** | **1.91e-15** |
| 48 | putative_ Chlorogenic_acid_ isomere | -7 | 1.9 | 0.16 | 1 | 6 | 0.02 | 0.9 | 1 | 44 | 95 | **2.2e-16** | **1.91e-15** |
| 49 | Glucose | 10 | 0.5 | 0.48 | 0.9 | 14 | 0.058 | 0.8 | 0.9 | -7 | 0.94 | 0.3 | 0.4 |
| 50 | Fructose | 19 | 0.95 | 0.33 | 0.9 | 17 | 0.27 | 0.6 | 0.9 | -14 | 2.27 | 0.13 | 0.2 |
| 51 | Sucrose | -1 | 0.12 | 0.72 | 0.7 | 10 | 1.1 | 0.3 | 0.7 | -21 | 10.8 | **0.001** | **0.0045** |

| | | | | | | | | | | | | | |
|---|---|---|---|---|---|---|---|---|---|---|---|---|---|
| 52 | Starch_(Glu) | -1 | 0.1 | 0.75 | 0.7 | 27 | 1.8 | 0.2 | 0.7 | -13 | 7.7 | **0.0056** | **0.0171** |

NA= no peak detected; *n=3 as for some replicates no peak detected, or peak too small for accurate quantification

**Table S6 Statistics for amplicon metagenomics of duckweed samples of outdoor mesocosms.** PERMANOVA testing the effect of treatment (herbivory/control) week of experiment (timepoint) on the structure of bacterial (16S) and algal (18S) communities associated with *Spirodela polyrhiza* material (with potentially adherent microalgae).

| **Factor** | | **16S (cyano-)bacteria** | | | **18S algae** | | |
|---|---|---|---|---|---|---|---|
| | **df** | **$R^2$** | **F** | ***p*-value** | **$R^2$** | **F** | ***p*-value** |
| *treatment* | 1 | 0.04 | 1.2 | 0.3 | 0.04 | 1.4 | 0.2 |
| *timepoint* | 2 | 0.19 | 3.1 | **0.001** | 0.18 | 3.1 | **0.001** |
| *treatment * timepoint* | 2 | 0.04 | 0.7 | 1 | 0.03 | 0.5 | 1 |

**Table S7 MRM-settings and retention times of analytes of Method 1A (metabolite analysis).**

| Analyte | RT [min] | Q1 [m/z] → | Q3 [m/z] | Dwell time [ms] | CE [V] | Q1/Q3 Pre Bias [V] | ISTD[a] |
|---|---|---|---|---|---|---|---|
| Ala | 0.450 | (+) 90,05 | 44,20 | 20 | -13 | -10 / -18 | $^{13}C_3$, $^{15}N_1$-Ala |
| Arg | 0.420 | (+)175,12 | 60,20 | 20 | -14 | -20 / -24 | $^{13}C_6$, $^{15}N_4$-Arg |
| Asp | 0.450 | (+)134,04 | 88,20 | 20 | -12 | -13 / -16 | $^{13}C_4$, $^{15}N_n$-$Asx_{Asp}$[b] |
| Glu | 0.450 | (+)148,06 | 102,15 | 20 | -13 | -10 / -19 | $^{13}C_5$, $^{15}N_n$-$Glx_{Glu}$[b] |
| Pro | 0.485 | (+)116,07 | 70,20 | 20 | -17 | -11 / -20 | $^{13}C_5$, $^{15}N_1$-Pro |
| Ser | 0.435 | (+)106,05 | 60,20 | 20 | -14 | -10 / -23 | $^{13}C_3$, $^{15}N_1$-Ser |
| Thr | 0.450 | (+)120,07 | 74,20 | 20 | -12 | -12 / -30 | $^{13}C_4$, $^{15}N_1$-Thr |
| Asn | 0.435 | (+)133,06 | 87,20 | 20 | -11 | -13 / -16 | $^{13}C_4$, $^{15}N_n$-$Asx_{Asn}$[b] |
| Gln | 0.450 | (+)147,08 | 130,15 | 20 | -15 | -10 / -13 | $^{13}C_5$, $^{15}N_n$-$Glx_{Gln}$[b] |
| Val | 0.627 | (+)118,09 | 72,20 | 20 | -12 | -22 / -28 | $^{13}C_5$, $^{15}N_1$-Val |
| Ile | 1.165 | (+)132,10 | 86,15 | 122 | -11 | -20 / -20 | $^{13}C_6$, $^{15}N_1$-Ile |
| Leu | 1.255 | (+)132,10 | 86,15 | 122 | -11 | -20 / -20 | $^{13}C_6$, $^{15}N_1$-Leu |
| Phe | 2.520 | (+)166,09 | 120,20 | 247 | -15 | -20 / -20 | $^{13}C_9$, $^{15}N_1$-Phe |
| Trp | 3.260 | (+)205,10 | 188,10 | 32 | -10 | -20 / -20 | $^{13}C_9$, $^{15}N_1$-Phe |
| (1.76) | | | | | | | |
| | | (+)205,10 | 118,20 | 32 | -26 | -20 / -20 | |
| $^{13}C_3$, $^{15}N_1$-Ala | 0.450 | (+) 94,06 | 47,20 | 20 | -13 | -10 / -18 | |
| $^{13}C_6$, $^{15}N_4$-Arg | 0.438 | (+)185,13 | 64,15 | 20 | -14 | -20 / -24 | |
| $^{13}C_4$, $^{15}N_n$-Asx | 0.450 | (+)139,06 | 92,25 | 20 | -12 | -13 / -16 | |
| $^{13}C_5$, $^{15}N_n$-Glx | 0.450 | (+)154,07 | 107,20 | 20 | -13 | -10 / -19 | |
| $^{13}C_5$, $^{15}N_1$-Pro | 0.485 | (+)122,08 | 75,20 | 20 | -17 | -11 / -20 | |
| $^{13}C_3$, $^{15}N_1$-Ser | 0.435 | (+)110,06 | 63,20 | 20 | -14 | -10 / -23 | |
| $^{13}C_4$, $^{15}N_1$-Thr | 0.450 | (+)125,08 | 78,20 | 20 | -12 | -12 / -30 | |
| $^{13}C_5$, $^{15}N_1$-Val | 0.627 | (+)124,10 | 77,20 | 20 | -12 | -22 / -28 | |
| $^{13}C_6$, $^{15}N_1$-Ile | 1.165 | (+)139,12 | 92,25 | 122 | -11 | -20 / -20 | |
| $^{13}C_6$, $^{15}N_1$-Leu | 1.255 | (+)139,12 | 92,25 | 122 | -11 | -20 / -20 | |
| $^{13}C_9$, $^{15}N_1$-Phe | 2.520 | (+)176,11 | 129,25 | 247 | -15 | -20 / -20 | |

RT: retention time

CE: collision energy

ISTD: internal standard

Qualifiers are depicted in grey

[a] Incl. matrix corrected response factor ($n_{Analyte} = x * n_{ISTD}$) in brackets

[b] Asx and Glx handled as Asn, Asp, Glu or Gln equivalents, respectively

**Table S8 MRM-settings and retention times of analytes of Method 1B (metabolite analysis).**

| Analyte | RT [min] | Q1 [m/z] | → | Q3 [m/z] | Dwell time [ms] | CE [V] | Q1/Q3 Pre Bias [V] | ISTD[a] |
|---|---|---|---|---|---|---|---|---|
| Gly | 0.460 | (+) 76,04 | | 30,20 | 50 | -12 | -14 / -11 | $^{13}C_2$, $^{15}N_1$-Gly |
| His | 0.440 | (+)156,08 | | 110,20 | 50 | -15 | -30 / -21 | $^{13}C_6$, $^{15}N_3$-His |
| Met | 0.750 | (+)150,06 | | 61,15 | 50 | -22 | -10 / -24 | $^{13}C_5$, $^{15}N_1$-Met |
| Shikimic acid (37.77) | 0.630 | (-) 173,05 | | 93,10 | 23 | 15 | 12 / 21 | $^{13}C_9$, $^{15}N_1$-Tyr |
| | | (-) 173,05 | | 111,20 | 23 | 13 | 12 / 10 | |
| Tyr | 1.130 | (+)182,08 | | 136,20 | 50 | -15 | -18 / -14 | $^{13}C_9$, $^{15}N_1$-Tyr |
| Tyramine (0.51) | 1.160 | (+)138,09 | | 121,15 | 23 | -15 | -20 / -20 | $^{13}C_9$, $^{15}N_1$-Tyr |
| | | (+)138,09 | | 77,15 | 23 | -30 | -20 / -20 | |
| Tryptamine (0.36) | 3.280 | (+)161,15 | | 144,15 | 297 | -15 | -20 / -20 | $^{13}C_9$, $^{15}N_1$-Tyr |
| | | (+)161,15 | | 117,15 | 297 | -25 | -20 / -22 | |
| Apigenin (0.96) | 4.120 | (-)269,05 | | 151,20 | 147 | 25 | 30 / 14 | $^{13}C_9$, $^{15}N_1$-Tyr |
| | | (-)269,05 | | 149,15 | 147 | 24 | 29 / 28 | |
| Luteolin (3.44) | 4.025 | (-)285,04 | | 132,20 | 147 | 48 | 30 / 26 | $^{13}C_9$, $^{15}N_1$-Tyr |
| | | (-)285,04 | | 175,20 | 147 | 26 | 19 / 11 | |
| $^{13}C_2$, $^{15}N_1$-Gly | 0.460 | (+) 79,04 | | 32,10 | 50 | -12 | -14 / -11 | |
| $^{13}C_6$, $^{15}N_3$-His | 0.440 | (+)165,09 | | 118,20 | 50 | -15 | -30 / -21 | |
| $^{13}C_5$, $^{15}N_1$-Met | 0.750 | (+)156,07 | | 63,15 | 50 | -22 | -10 / -24 | |
| $^{13}C_9$, $^{15}N_1$-Tyr | 1.130 | (+)192,11 | | 145,20 | 50 | -15 | -18 / -14 | |

RT: retention time

CE: collision energy

ISTD: internal standard

Qualifiers are depicted in grey

[a] Incl. matrix corrected response factor ($n_{Analyte} = x * n_{ISTD}$) in brackets

**Table S9 MRM-settings and retention times of analytes of Method 1C (metabolite analysis).**

| Analyte | RT [min] | Q1 [m/z] | → | Q3 [m/z] | Dwell time [ms] | CE [V] | Q1/Q3 Pre Bias [V] | ISTD[a] |
|---|---|---|---|---|---|---|---|---|
| Fructose | 7.8 | (-)179,06 | | 88,90 | 19 | 9 | 12 / 13 | Sorbitol[a] |
| | | (-)179,06 | | 59,00 | 19 | 17 | 12 / 11 | |
| | | (-)179,06 | | 71,00 | 19 | 16 | 11 / 11 | |
| Glucose | 9.2 | (-)179,06 | | 88,90 | 19 | 9 | 12 / 13 | Sorbitol[a] |
| | | (-)179,06 | | 59,00 | 19 | 17 | 12 / 11 | |
| | | (-)179,06 | | 71,00 | 19 | 16 | 11 / 11 | |
| Sucrose | 8.4 | (-)341,11 | | 89,10 | 19 | 23 | 12 / 10 | Sorbitol[a] |
| | | (-)341,11 | | 179,15 | 19 | 15 | 30 / 30 | |
| | | (-)341,11 | | 59,05 | 19 | 36 | 29 / 23 | |
| Sorbitol | 10.8 | (-)181,07 | | 88,85 | 19 | 16 | 11 / 17 | |
| | | (-)181,07 | | 59,00 | 19 | 22 | 11 / 10 | |
| | | (-)181,07 | | 71,00 | 19 | 22 | 30 / 29 | |

RT: retention time

CE: collision energy

ISTD: internal standard

Qualifiers are depicted in grey

[a] Response factor calculated based on an external dilution curve run with each batch of samples

**Table S10 Absorption and retention times of analytes of Method 1D (metabolite analysis).**

| Analyte | RT [min] | Absorption wave lenght [nm] |
|---|---|---|
| Cyanidin-3-O-glucoside | 7.055 | 517 |
| Chlorogenic acid | 8.055 | 328 |
| Putative Chlorogenic acid isomere | 8.210 | 328 |
| Cyanidin-3-O-(6-O-malonyl-beta-glucoside) | 9.410 | 517 |
| Luteolin-8-C glucoside | 11.295 | 348 |
| Apigenin-8-C-glucoside | 12.495 | 337 |
| Luteolin-7-O-glucoside | 13.085 | 348 |
| Apigenin-7-O-glucoside | 14.325 | 337 |

RT: retention time

Quantification based on an external dilution curve with identical standards, except for the putative chlorogenic acid isomere that was based on the chlorogenic acid standard, and cyanidin-3-O-(6-O-malonyl-beta-glucoside) that was quantified based on the molar quantity of the cyanidin-3-O-glucoside standard.

**Table S11 MRM-settings and retention times of analytes of Method 2A (metabolite analysis).**

| Analyte | RT [min] | Q1 [m/z] | → | Q3 [m/z] | Dwell time [ms] | CE [V] | Q1/Q3 Pre Bias [V] | ISTD[a] |
|---|---|---|---|---|---|---|---|---|
| SA | 2.570 | (-)137,02 | | 92,95 | 20 | 19 | 27 / 19 | $D_4$-SA |
| | | (-)137,02 | | 65,00 | 20 | 32 | 27 / 10 | |
| ABA | 2.710 | (-)263,13 | | 153,10 | 20 | 13 | 18 / 29 | $D_6$-ABA |
| | | (-)263,13 | | 204,15 | 20 | 20 | 18 / 12 | |
| JA | 3.200 | (-)209,12 | | 59,00 | 20 | 14 | 14 / 11 | $D_5$-JA |
| | | (-)209,12 | | 40,95 | 20 | 44 | 14 / 14 | |
| JA-Ile | 4.570 | (-)322,20 | | 130,10 | 135 | 22 | 24 / 20 | $D_5$-JA (0.084) |
| | | (-)322,20 | | 128,15 | 135 | 22 | 24 / 26 | |
| OPDA | 6.210 | (-)291,20 | | 165,25 | 135 | 22 | 20 / 10 | $D_5$-JA[b] |
| | | (-)291,20 | | 247,15 | 135 | 20 | 20 / 23 | |
| $D_4$-SA | 2.560 | (-)141,05 | | 97,00 | 20 | 19 | 27 / 15 | |
| | | (-)141,05 | | 69,05 | 20 | 32 | 27 / 17 | |
| $D_6$-ABA | 2.700 | (-)269,17 | | 159,20 | 20 | 13 | 18 / 30 | |
| | | (-)269,17 | | 207,20 | 20 | 20 | 18 / 20 | |
| $D_5$-JA | 3.190 | (-)214,15 | | 62,00 | 20 | 14 | 14/ 10 | |
| | | (-)214,15 | | 42,00 | 20 | 44 | 14 / 15 | |

RT: retention time

CE: collision energy

ISTD: internal standard

Qualifiers are depicted in grey

[a] Incl. matrix and recovery corrected response factor ($n_{Analyte} = x * n_{ISTD}$) in brackets

[b] Relative quantification

**Table S12 MRM-settings and retention times of analytes of Method 2B (metabolite analysis).**

| Analyte | RT [min] | Q1 [m/z] → | Q3 [m/z] | Dwell time [ms] | CE [V] | Q1/Q3 Pre Bias [V] | ISTD[a] |
|---|---|---|---|---|---|---|---|
| Caffeic acid | 2.275 | (-) 179,03 | 135,05 | 20 | 18 | 12 / 26 | 4-MU[b] |
| | | (-) 179,03 | 134,15 | 20 | 25 | 12 / 12 | |
| | | (-) 179,03 | 107,10 | 20 | 24 | 19 / 21 | |
| *p*-Coumaric acid | 2.624 | (-) 163,04 | 119,10 | 20 | 17 | 11 / 11 | 4-MU[b] |
| | | (-) 163,04 | 93,05 | 20 | 32 | 11 / 19 | |
| Sinapic acid | 2.760 | (+)225,08 | 207,15 | 20 | -9 | -15 / -23 | 4-MU[b] |
| | | (+)225,08 | 119,20 | 20 | -19 | -15 / -12 | |
| IAA | 3.237 | (+)176,07 | 130,10 | 20 | -16 | -12 / -23 | $^{13}C_6$-IAA |
| (0.42) | | | | | | | |
| | | (+)176,07 | 103,20 | 20 | -31 | -12 / -10 | |
| Cinamic acid | 5.002 | (-) 147,05 | 103,10 | 100 | 15 | 17 / 20 | 4-MU[b] |
| | | (-) 147,05 | 77,10 | 100 | 23 | 10 / 18 | |
| $^{13}C_6$-IAA | 3.237 | (+)182,09 | 109,20 | 20 | -31 | -12 / -19 | |
| | | (+)182,09 | 136,25 | 20 | -16 | -12 / -13 | |
| 4-MU | 3.344 | (+)177,05 | 77,20 | 20 | -35 | -12 / -30 | |
| | | (+)177,05 | 105,20 | 20 | -21 | -12 / -20 | |

RT: retention time

CE: collision energy

ISTD: internal standard

Qualifiers are depicted in grey

[a] Incl. response factor ($n_{Analyte} = x * n_{ISTD}$) in brackets

[b] Relative quantification

**Table S13 MRM-settings and retention times of analytes of Method 3 (metabolite analysis).**

| Analyte | RT [min] | Q1 [m/z] → | Q3 [m/z] | Dwell time [ms] | CE [V] | Q1/Q3 Pre Bias [V] | ISTD[a] |
|---|---|---|---|---|---|---|---|
| tZ | 2.527 | (+)220,12 | 136,10 | 47 | -20 | -20 / -20 | $D_5$-tZ |
| | | (+)220,12 | 119,15 | 47 | -33 | -22 / -12 | |
| cZ | 2.830 | (+)220,12 | 136,10 | 47 | -20 | -20 / -20 | $D_5$-tZ (0.94) |
| | | (+)220,12 | 119,15 | 47 | -33 | -22 / -12 | |
| tZR | 4.292 | (+)352,16 | 220,15 | 30 | -20 | -20 / -20 | $D_5$-tZ (0.17) |
| | | (+)352,16 | 136,15 | 30 | -30 | -20 / -20 | |
| cZR | 4.696 | (+)352,16 | 220,15 | 30 | -20 | -20 / -20 | $D_5$-tZ (0.17) |
| | | (+)352,16 | 136,15 | 30 | -33 | -20 / -20 | |
| IP | 5.648 | (+)204,12 | 136,10 | 30 | -15 | -20 / -20 | $D_6$-IP |
| | | (+)204,12 | 119,10 | 30 | -30 | -20 / -20 | |
| IPR | 6.390 | (+)336,17 | 204,20 | 47 | -20 | -20 / -20 | $D_6$-IPR |
| | | (+)336,17 | 136,10 | 47 | -30 | -20 / -20 | |

| | | | | | | |
|---|---|---|---|---|---|---|
| $D_5$-tZ | 2.520 | (+)225,15 | 137,20 | 47 | -20 | -21 / -22 |
| | | (+)225,15 | 119,20 | 47 | -36 | -15 / -12 |
| $D_6$-IP | 5.511 | (+)210,16 | 137,20 | 30 | -15 | -20 / -20 |
| | | (+)210,16 | 119,15 | 30 | -33 | -14 / -22 |
| $D_6$-IPR | 6.363 | (+)342,20 | 210,25 | 47 | -20 | -20 / -20 |
| | | (+)342,20 | 137,15 | 47 | -30 | -20 / -20 |

RT: retention time

CE: collision energy

ISTD: internal standard

Qualifiers are depicted in grey

[a] Incl. matrix and recovery corrected response factor ($n_{Analyte} = x * n_{ISTD}$) in brackets

**Table S14 Solvent settings used for Method 1A**

| Time [min] | B [%] |
|---|---|
| *0,0* | *2* |
| *1,5* | *2* |
| *3,5* | *100* |
| *4,5* | *100* |
| *5,0* | *2* |
| *6,0* | *2* |

**Table S15 Solvent settings used for Method 1B**

| Time [min] | B [%] |
|---|---|
| *0,0* | *2* |
| *1,5* | *2* |
| *3,5* | *100* |
| *4,5* | *100* |
| *5,0* | *2* |
| *6,0* | *2* |

**Table S16 Solvent settings used for Method 1C**

| Time [min] | B [%] |
|---|---|
| *0* | *80* |
| *13* | *60* |
| *14* | *80* |
| *20* | *80* |

**Table S17 Solvent settings used for Method 1D**

| Time [min] | B [%] |
|---|---|
| *0,0* | *10* |
| *8,0* | *21* |
| *18,0* | *49* |
| *18,1* | *100* |
| *19,0* | *100* |
| *19,1* | *10* |
| *24,0* | *10* |

**Table S18 Solvent settings used for Method 2A**

| **Time [min]** | **B [%]** |
|---|---|
| *0,0* | *10* |
| *0,5* | *10* |
| *1,0* | *55* |
| *4,5* | *65* |
| *5,5* | *100* |
| *6,5* | *100* |
| *7,0* | *10* |
| *8,0* | *10* |

### Table S19 Solvent settings used for Method 2B

| Time [min] | B [%] |
|---|---|
| *0,0* | *10* |
| *0,5* | *10* |
| *1,0* | *39* |
| *3,5* | *41* |
| *3,6* | *50* |
| *8,0* | *60* |
| *8,5* | *100* |
| *9,5* | *100* |
| *10,0* | *10* |
| *11,0* | *10* |

### Table S20 Solvent settings used for Method 3

| Time [min] | B [%] |
|---|---|
| *0,0* | *5* |
| *0,5* | *5* |
| *0,7* | *15* |
| *3,5* | *25* |
| *6,5* | *70* |
| *6,7* | *100* |
| *7,7* | *100* |
| *8,0* | *5* |
| *9,0* | *5* |

### TEXT S1 DETAILED METHODS on cultivation and preadaptation of species

To gain starting material for both indoor and outdoor experiments, *Spirodela polyrhiza* was cultivated in the growth chamber in 1 l flasks in liquid full N-Medium [4] under sterile conditions for four weeks. Cultures of the microalgae *Chlamydomonas reinhardii* were cultivated in TAP-Medium (SI Table S1, based on (Hutner et al., 1950)) within the same chamber for two weeks. Population of *Lymnaea stagnalis* was cultivated at room temperature in aquariums in our lab and fed with frozen spinach and commercial fish feed.

Prior indoor experiments, *S. polyrhiza* was always transferred to Mix-Medium (SI Table S2) to preadapt to experimental conditions for seven days. Algae cultures were always preadapted in Mix-Medium for 24 h, using gentle centrifugation to exchange the media. Snails were always starved for 24 h within Mix-Medium in the same growth chamber in which experiments were conducted. Snails had a shell size of 14-32 mm and we used groups of similar size range across treatments.

Our approach is based on pretests testing the overall feasibility of our indoor microcosm set-up, e. g. by testing the maxium capacity of *S. poylrhiza* fronds for given space, testing density gradients of plants and algae, and given nutrient levels. Further we performed tests to ensure “ad libitum” conditions when quantifying consumption rates of duckweed and algae when testing consumption rates.

### TEXT S2 DETAILED METHODS for laborartory 3-species microcosm setup

We here aimed to test within laboratory microcosms under which starting conditions *Spirodela polyrhiza* (*Sp*) may benefit from the presence of snail herbivory when competing with *C. reinhardii* (*Ch*), to then comare the outcome with our theoretic model. We excluded those replicates from analysis, in which duckwed populations collabsed or snails died within 14 days of the experiment, resulting in the below-stated replicate numbers. All treatments were further set up in the presence of two snails. Initially we used n=4 for all treatments. As when using two snails many snails died along the two weeks, we fully excluded this data set from analysis but kept the corresponding controls, leading to n=8.

- *Sp* high (300 fronds) + *Ch* low (0.9 cells [ml ($x^{*10^\wedge 6}$)]$^{-1}$) without 1 snail (n=4)
- *Sp* high (300 fronds) + *Ch* low (0.9 cells [ml ($x^{*10^\wedge 6}$)]$^{-1}$) with 1 snail (n=4)

- *Sp* medium (200 fronds) + *Ch* medium-low (1.3 cells [ml ($x^{*10^\wedge 6}$)]$^{-1}$) without 1 snail (n=8)

- *Sp* medium (200 fronds) + *Ch* medium-low (1.3 cells [ml ($x^{*10^\wedge 6}$)]$^{-1}$) with 1 snail (n=3)

- *Sp* medium (200 fronds) + *Ch* medium-low (4.2 cells [ml ($x^{*10^\wedge 6}$)]$^{-1}$) without 1 snail (n=8)
- *Sp* medium (200 fronds) + *Ch* medium-low (4.2 cells [ml ($x^{*10^\wedge 6}$)]$^{-1}$) with 1 snail (n=3)

- *Sp* low (100 fronds) + *Ch* high (5.8 cells [ml ($x^{*10^\wedge 6}$)]$^{-1}$) without 1 snail (n=4)
- *Sp* low (100 fronds) + *Ch* high (5.8 cells [ml ($x^{*10^\wedge 6}$)]$^{-1}$) with 1 snail (n=3)

### TEXT S3 DETAILED METHODS to calculate surface and biomass of *S. polyrhiza* grown in outdoor mesocosms

Normalized fitness change was calculated by using the coverage rates in percent with the following formula: (Herbivory – Control / Maximum value) for each timepoint, respectively. Additionally, we sampled 20 fronds with roots randomly picked from each treatment zone per pond in week 0, 2, 3, 4, 5, 7, 8 and 9 and measured biomass.

The effect of herbivory on plant fitness measured via surface area coverage and biomass was tested by fitting a linear-mixed effect model, using coverage or biomass within each treatment zone as variable and specifying *treatment* (herbivory/control) as fixed factor, and *pond* and *timepoint* as random factors, using the formula: ~ treatment * (1|week) * (1|pond).

### TEXT S4 DETAILED METHODS for metabolite analysis of duckweed grown in outdoor mesocosms

To test whether multigenerational herbivory outdoors leads to changes in the production of a wide set of plant metabolites, using HPLC and LC-MS analysis, we collected a set of 30 fronds randomly picked from each treatment from each pond at week 0, 3 and 6 (n=5 per timepoint for week 0 and 3, n=3 for week 6 due to sample loss). Samples were freeze-dried prior extraction.

We tested the effect of herbivory (herbivory/control) for each metabolite by fitting a linear-mixed effect model, using the concentration within each treatment zone as variable and specifying *treatment* (herbivory/control) as fixed factor, and *pond* and *timepoint* as random factors, using the formula: ~ treatment * (1|week) * (1|pond).

Extraction and analysis of diverse primary and secondary metabolites was done similar as described by [5]. A detailed list of the analysed compound and the respective analytical methods can be found in SI Table S7-13. In brief, approximately 10 mg fine ground, freeze dried plant tissue was extracted with acidified methanol containing the following internal standards: 10 ng $D_6$-ABA (OlChemIm); 10 ng $D_5$-JA (Sigma-Aldrich); 10 ng $D_4$-SA (OlChemIm); 0.5 ng $D_5$-tZ (OlChemIm); 0.1 ng $D_6$-IP (OlChemIm); 0.1 ng $D_6$-IPR (OlChemIm); 1 ng $^{13}C_6$-IAA (OlChemIm); 3 ng 4-methylumbelliferone (4-MU) (Sigma-Aldrich). For the analysis of different high abundant compounds, a small aliquot (A1) of the extract was taken aside, while the rest of the extract (E1) was used for further processing. High abundant compounds, such as amino acids and related compounds were analysed in a 1:100 dilution of A1 in an aqueous mix of isotope-labeled amino acids (algal amino acid mixture-$^{13}C$-$^{15}N$; Sigma-Aldrich; Method 1A and 1B). Soluble sugars were measured in a 1:125 dilution of A1 in 70 % methanol containing sorbitol as internal standard (Fluka Biochemika; Method 1C). The major secondary metabolites were analysed in the rest of the aliquot (A1) without further dilution (Method 1D). After a second extraction of the plant tissue with acidified methanol this supernatant was combined with the first extract (E1) and subsequently purified on a reversed phase (RP) solid phase extraction (SPE) column (HR-X, Macherey-Nagel) and a mixed-mode RP-cation exchange SPE column (HR-XC, Macherey-Nagel). The compounds were bound to the HR-XC column, different washing steps were applied (e.g. to remove CK-phosphates) and the analytes of interest were stepwise eluted from the column to generate a fraction containing the acidic phytohormones, such as JA, ABA and IAA, as well as a fraction containing the CKs. The eluted fractions were analysed directly (Method 2A) or after an additional concentration step (Method 2B) and (Method 3) dependent on the abundance and instrument sensitivity for the respective compounds. Starch content was determined from the remaining material (pellet) after the two rounds of extraction with acidified methanol. For this, the material was subjected to two additional washing steps with 50 % ethanol at 80 °C to remove soluble sugars. Subsequently, a sub-fraction of the material was resuspended in water and incubated at 98 °C to gelatinize the starch, which was then digested to glucose by amyloglucosidase (from *Aspergillus niger*, Roche) and αamylase (from porcine pancreas, Sigma) at 37 °C. The glucose content was then measured in a 1:250 dilution of this solution in 70 % methanol containing sorbitol as internal standard the same way as for the soluble sugars before (Method 1C).

The major secondary metabolites (Method 1D) were analysed by HPLC-PDA. Chromatographic separation was done on a Shimadzu Nexera XR LC-System equipped with an EC 4/3 Nucleodur Sphinx RP pre-column (5 µm, Macherey-Nagel) and a Nucleodur Sphinx RP column (250x4.6 mm, 5µm, Macherey-Nagel). The mobile phase comprised 0.2 % formic acid (Fisher Chemical), 0.1 % acetonitrile (Fisher Chemical) in water as Solvent A and acetonitrile (Fisher Chemical) as Solvent B. The column oven was set to 20 °C and the flow rate to 1300 µL/min. Measurement was performed with a PDA detector.

Other compounds (Method 1A, 1B, 1C, 2A, 2B and 3) were analysed by LC-MS/MS. Chromatographic separation for was done on a Shimadzu Nexera X3 LC-System. For method 1A, 1B, 2A, 2B and 3 the system was equipped with an Agilent 1290 infinity II inline filter (0,3 µm) and a ZORBAX RRHD Eclipse XDB-C18 column (3x50 mm, 1.8 µm; Agilent Technologies). The mobile phase comprised 0.05 % formic acid (Fisher Chemical), 0.1 % acetonitrile (Fisher Chemical) in water as Solvent A and methanol (Fisher Chemical) as Solvent B. The column oven was set to 42 °C and the flow rate to 500 µL/min. For method 1C the system was equipped with an Agilent 1290 infinity II inline filter (0,3 µm) and an apHera NH2 column (150x4.6 mm, 5 µm; Supelco). The mobile phase comprised 0.1 % acetonitrile (Fisher Chemical) in water as Solvent A and acetonitrile (Fisher Chemical) as Solvent B. The column oven was set to 25 °C and the flow rate to 700 µL/min. The measurements were performed on a Shimadzu LCMS-8060 equipped with an ESI source, which was operated in multi-reaction-monitoring (MRM) modus. Settings were as follows: Nebulizing Gas Flow: 3 L/min; Heating Gas Flow: 10 L/min; Drying Gas Flow: 10 L/min; Interface Temperature: 300 °C; DL Temperature 250 °C; Heat Block Temperature: 400 °C; CID Gas: 270 kPa; Q1 Resolution: Unit; Q3 Resolution: Unit. The gradient programs for all methods are given in SI Table S14-20 and the detector settings (MRM-settings or absorption wavelength, respectively) in SI Table S7-13.

**TEXT S5 DETAILED METHODS for water analysis of outdoor mesocosms.**

To test for nutrient levels between compartment zones of outdoor ponds, we analysed nutrients using ion-chromatography. Samples were collected in week 0, 3, 5, 7 and 9. We sampled 10 ml per treatment zone for each anion and cation analysis. Water was collected from two locations within each zone at a depth of 10 cm below the water surface. Anion (Cl, $NO_2$, $NO_3$, $PO_4$, $SO_4$) samples were measured right away, while pH of cation samples (for Na, $NH_4$, K, Ca, Na) was adjusted to pH 3 using 25 % $HNO_3$ before storage at -20°C until analysis.

The effect of herbivory on water nutrients was tested by fitting a linear-mixed effects model specifying *treatment* (herbivory/control) as fixed factor, and *pond* and *timepoint* as random effects, using the formula: ~ treatment * (1|timepoint) * (1|pond). In addition, we tested the effects of overall nutrient content through PERMANOVA (999 permutations, stratified over "*pond*" and "*sampling timepoint*").

Water temperature in the reach of duckweed was measured using a data logger (HoboPendant MXTemp, *Onset Computer Corporation*, MA, USA*)*. Data for air temperature was provided by a weather station about 1 km away from site.

### REFERENCES


1. Stevens H. 2021 Primer of Ecology using R. See https://hankstevens.github.io/Primer-of-Ecology/comp.html (accessed on 5 February 2023).
2. Hutner SH, Provasoli L, Schatz A, Haskins CP. 1950 Some Approaches to the Study of the Role of Metals in the Metabolism of Microorganisms. *Proc Am Philos Soc* **94**, 152–170.
3. Hutner SH, Provasoli L, Schatz A, Haskins CP. 1950 Some Approaches to the Study of the Role of Metals in the Metabolism of Microorganisms. *Proc Am Philos Soc* **94**, 152–170.
4. Appenroth KJ, Teller S, Horn M. 1996 Photophysiology of turion formation and germination in Spirodela polyrhiza. *Biol Plant* **38**, 95–106. (doi:10.1007/BF02879642)
5. Schäfer M, Brütting C, Baldwin IT, Kallenbach M. 2016 High-throughput quantification of more than 100 primary- and secondary-metabolites, and phytohormones by a single solid-phase extraction based sample preparation with analysis by UHPLC-HESI-MS/MS. *Plant Methods* **12**. (doi:10.1186/s13007-016-0130-x)
